# *Phytolacca americana Pa*GT2 is an ambidextrous polyphenol glucosyltransferase

**DOI:** 10.1101/620955

**Authors:** Rakesh Maharjan, Yohta Fukuda, Naomichi Shimomura, Taisuke Nakayama, Toru Nakayama, Hiroki Hamada, Tsuyoshi Inoue, Shin-ichi Ozaki

## Abstract

The health benefits of polyphenols have attracted their use as potential therapeutic agents, food additives, and cosmetics. However, low water solubility of polyphenols limits their cell absorbability, obscuring further exploration. Glycosylation is known to enhance the solubility of polyphenols preserving their pharmacological properties. Here, we show that a uridine diphosphate (UDP) glucosyltransferase from *Phytolacca americana* (*Pa*GT2) regioselectively catalyzes the transfer of glucose from UDP-glucose to stilbenoids such as piceatannol and flavonoids such as kaempferol. To understand the structure-function relationship of *Pa*GT2, we determined the crystal structure of *Pa*GT2 as well as *Pa*GT2 complexed with donor analogue UDP-2-fluoro glucose and stilbenoid acceptor analogues. While only one conserved histidine residue is recognized as a catalytic residue in known UGTs, the crystal structures of *Pa*GT2 suggested the presence of two catalytically active residues (His18 and His81) at two sides of the catalytic pocket. Although the single catalytic residue mutant His18Ala or His81Ala did not completely impair the glucosylation activity of the enzyme, the double mutant His18Ala/His81Ala failed to form glucoside products. These results showed that both catalytic residues in *Pa*GT2 actively and independently catalyze glucosylation, hence we called *Pa*GT2 as an ambidextrous UGT. The information from *Pa*GT2 will be advantageous for the engineering of efficient biocatalysts for production of therapeutic polyphenols.

## Introduction

Glycosylation of secondary metabolites is one of the important mechanism in plants for their metabolism, intercellular/intracellular localization, and storage (1). Glycosylation of xenobiotic compounds also play a major role in detoxification system of plants and thereby minimize the risk of toxic components from the environment (2). Thus, glycosylation of lipophilic molecules is one of the essential mechanisms for maintaining cellular homeostasis (3). In plants, the addition of sugar moieties to small molecules is catalyzed by uridine diphosphate glycosyltransferases (UGTs) which are classified as GT1 family in Carbohydrate Active Enzyme (CAZy) database (4, 5). Enzymes in this family show catalytic plasticity; that is, they are able to glycosylate a wide variety of acceptors (9–11) as well as glycosylate a single acceptor at various available glycosylation sites (12, 13).

The UGTs in GT1 family have glycosyltransferase-B (GT-B) fold structures, made up of two Rossmann fold domains which are connected by a linker (6). Plant GT1 enzymes are characterized with the presence of consensus sequence known as plant secondary product glycosyltransferase (PSPG) motif that is involved in recognition of the UDP-sugar donor (7). The N-terminal domain of these UGTs has a highly conserved histidine which is considered to be the main catalytic residue (3, 8). The acceptor binding site at the N-terminal domain has low sequence similarity among UGTs, reflecting a wide spectrum of substrates. However, only the difference in acceptor binding residues among UGTs is not enough to explain the substrate plasticity of these enzymes.

Polyphenols such as stilbenoids and flavonoids are known to have antioxidant, anti-estrogenic, anticancer, anti-inflammatory, and cardio-protective effects (14–17), and have important applications in food, pharmaceuticals, and cosmetics industries (18). Most of these polyphenols are found to be glycosylated in plants (9, 19) and differences in positions of glycosylation sites can result in different biological activity of glycosides (17). Flavonoid glucosides such as kaempferol-3-*O*-glucoside and quercetin-3-*O*-glucoside have been identified in the leaves of *Phytolacca americana*, a toxic plant native to North America (20). From the roots of this plant, triterpene saponin glucosides have been extracted and characterized (21). Three glycosyltransferases namely *Pa*GT1, *Pa*GT2, and *Pa*GT3 have been isolated from the callus tissues of *P. americana* (11). Among the three *Pa*GTs, *Pa*GT3 can glycosylate a wide variety of substrates, such as capsaicin, flavonoids, hydroxyflavones, and stilbenoids (11, 22, 23). Although *Pa*GT2 can also regioselectively glucosylate stilbenoids and flavonoids, the detail has not been much explored.

A large number of plant UGT gene sequences have been deposited in database; however, crystal structures of only few of them are available. These crystal structures of UGTs were studied for the glycosylation of iso/flavonoids (3, 4, 24) or small molecules, such as trichlorophenol (25), and indoxyl sulfate (26). Consequently, crystal structures of plant UGT with stilbenoids are not available. Stilbenoids are more flexible polyphenols compared to flavonoids and the existing UGT structures would not be sufficient to understand the mechanism of stilbenoid glycosylation. Hence, to improve our understanding of glycosylation mechanism and plasticity, crystal structures of more UGTs complexed with their putative substrates should be determined. In this study we report the crystal structure of apo-*Pa*GT2 and *Pa*GT2 complexed with stilbenoids, such as resveratrol and ptrerostilbene along with sugar donor analogue UDP-2-fluoro glucose (UDP-2FGlc). The identification of key residues, mutational studies, and comparison with other UGT structures provide a basis for understanding the regioselective glucosylation of acceptors by *Pa*GT2.

## Results

### Screening of stilbenoids for glucosylation by *Pa*GT2

The glucosylation activity of *Pa*GT2 was determined with UDP-glucose as the glucose donor. Various stilbenoids and kaempferol (as a representative for flavonoids) were utilized as the acceptor substrates (Fig. 1, Table 1). Stilbenoids and flavonoids are structurally different polyphenols. The lack of aromatic ring that bridge ring A and ring B in stilbenoids makes them more flexible when compared with flavonoids. *Pa*GT2 transformed piceatannol into piceatannol 4’-*O*-β-glucoside but did not form pterostilbene and rhapointigenin glucosides despite having molecular frameworks similar to piceatannol (Figs. 1, S2). Even more surprisingly, *Pa*GT2 could glucosylate only trace amount of resveratrol that only lacks the 3’OH group compared to piceatannol. Kaempferol was converted to kaempferol 3-*O*-β-glucoside by *Pa*GT2 although the *k*_cat_ value was one quarter of that for piceatannol. The results indicated piceatannol as the preferred substrate for *Pa*GT2. Notably, piceatannol and kaempferol both have four possible glucosylation sites; however, only one site on each compound was glucosylated with high regioselectivity. While the 4’OH in piceatannol and kaempferol seems to be structurally equivalent, only piceatannol 4’-*O*-β-glucoside was formed. To elucidate the molecular basis for this intriguing reactivity of *Pa*GT2, we performed structural analysis as following.

**Fig. 1.**
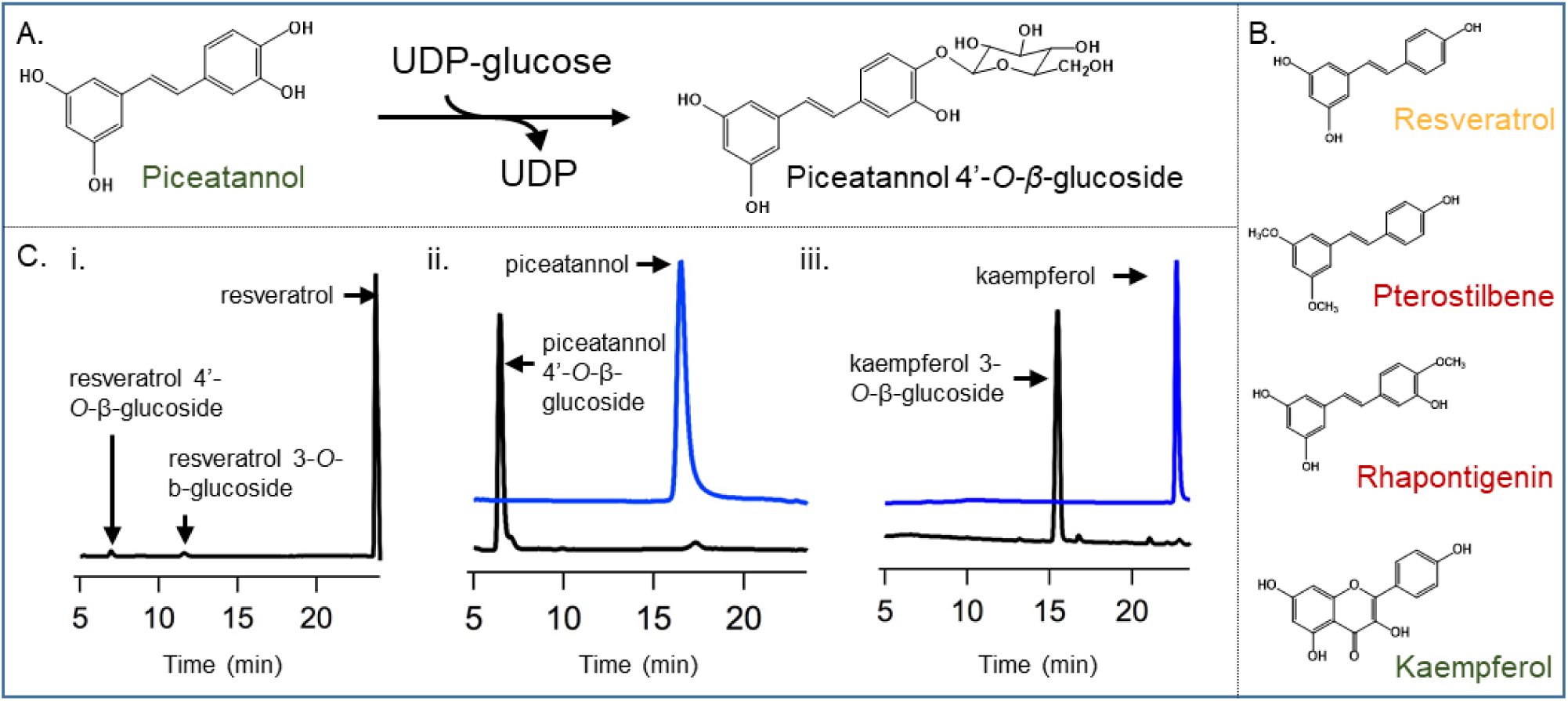
Panel of acceptors used for glucosylation study and HPLC analysis of products. (A) Schematic diagram of reaction catalyzed by *Pa*GT2 (B) Other compounds screened for glucosylation by *Pa*GT2. The names of aglycones are colored where green and orange mean good and poor acceptor, respectively, and red shows that the molecules were not catalyzed by *Pa*GT2. (C) HPLC analysis of glucosylation reaction catalyzed by *Pa*GT2 with UDP-glucose and substrates; i. resveratrol, ii. piceatannol, and iii. kaempferol. Peaks obtained for substrate and products in chromatograms are indicated. Blue and black lines show chromatograms before and after the reactions, respectively.

**Table 1:** Kinetic data of *Pa*GT2 and mutants

| Enzyme<br>(*PaGT2*) | Acceptor | $K_m$ ($\mu\text{M}$) | $k_{\text{cat}}$ ($\text{s}^{-1}$) | $k_{\text{cat}}/K_m$ ($\text{M}^{-1} \text{s}^{-1}$) |
| --- | --- | --- | --- | --- |
| WT | Piceatannol | $29 \pm 2^*$ | $1.46 \times 10^{-2} \pm 1.16 \times 10^{-3}$ | $505.74 \pm 53$ |
| H18A | | $20 \pm 3$ | $4.50 \times 10^{-3} \pm 6.67 \times 10^{-4}$ | $225.00 \pm 47$ |
| H81A | | $52 \pm 3$ | $8.66 \times 10^{-3} \pm 1.83 \times 10^{-3}$ | $166.66 \pm 36$ |
| E82A | | $40 \pm 3$ | $4.33 \times 10^{-3} \pm 1.16 \times 10^{-3}$ | $108.33 \pm 30$ |
| C142A | | $49 \pm 2$ | $2.16 \times 10^{-2} \pm 1.66 \times 10^{-3}$ | $442.17 \pm 38$ |
| H18A/H81A |  | N.D. | N.D. | N.D. |
| WT | Kaempferol† | $33 \pm 7$ | $3.66 \times 10^{-3} \pm 3.33 \times 10^{-4}$ | $111.11 \pm 25$ |
| H18A | | $26 \pm 4$ | $3.50 \times 10^{-3} \pm 1.67 \times 10^{-4}$ | $137.42 \pm 25$ |
| H81A | | $64 \pm 5$ | $6.16 \times 10^{-3} \pm 1.66 \times 10^{-3}$ | $96.35 \pm 27$ |
| E82A | | $40 \pm 2$ | $6.17 \times 10^{-3} \pm 1.36 \times 10^{-4}$ | $157.99 \pm 7$ |
| C142A | | $41 \pm 2$ | $5.16 \times 10^{-3} \pm 1.17 \times 10^{-4}$ | $127.65 \pm 4$ |
| H18A/H81A |  | N.D. | N.D. | N.D. |
| WT | Resveratrol | N.D. | N.D. | N.D. |
| WT | Pterostilbene | N.D. | N.D. | N.D. |
| WT | Rhapointigenin | N.D. | N.D. | N.D. |

\* Data are presented by means ± SEM (standard error of the mean, $n = 3$ ).
† Kinetic parameters for kaempferol determined for overall glucoside products. The H18A, E82A and C142A mutants produced a mixture of glucosides including kaempferol 3-*O*- β-glucoside.

### Structure of *Pa*GT2 without substrates

The crystal structure of *Pa*GT2 in its apo form was solved by molecular replacement using the structure of *Arabidopsis thaliana* UGT72B1 (PDB: 2VCH) (25) as a search model and refined to 2.30 Å resolution (Fig. S3, SI Table 1). The asymmetric unit contained two *Pa*GT2 molecules that were highly similar to each other with overall root mean square deviation (rmsd) of 0.29 Å for overlap of 366 Cα atoms. The structure of *Pa*GT2 belonged to GT-B fold, made up of two Rossmann (β/α/β) domains (27). The N-terminal domain (residues Ala5-Ser243) contained central seven parallel β-sheet flanked by eight α-helices which include an α-helix from C-terminal domain, and a small two-stranded β-sheet. The C-terminal domain (residues Ser252-Gln466) consisted of six β-sheet surrounded by eight α-helices. The loop (Gly244-Gly251) that connected the N-terminal domain with the C-terminal domain was disordered in the structure. *Pa*GT2 molecules in the asymmetric unit were dimerized by insertion of a long loop (Val300-Gly328) from the opposite molecules into the acceptor binding pockets (Fig. S3). The dimerization of *Pa*GT2 in the crystal structure was an artificial effect of crystallization (molecular packing, high concentration of protein, effect of precipitant, etc.) because the enzyme in solution showed a single symmetrical peak corresponding to its monomeric molecular weight in size exclusion chromatography (SEC) (Fig. S1). The structure of *Pa*GT2 also contained a kinked C-terminal helix that crossed over to N-terminal domain, a characteristic feature of GT-B fold UGT structures (28).

*Pa*GT2 showed the highest sequence similarity of 58% with UGT72B1 (25) and 56% with *Pt*UGT1 from *Polygonum tinctorium* (26). The overall structure of *Pa*GT2 was also highly similar with the structures of UGT72B1 and *Pt*UGT1 (PDB: 5NLM) with rmsds of 1.16 Å for 371 Cα atoms and 1.03 Å for 373 Cα atoms, respectively (Fig. S5A).

### Structure of *Pa*GT2 with substrates

*Pa*GT2 was crystallized with donor analogue UDP-2-fluoro-glucose (UDP-2FGlc) and various acceptors: resveratrol, pterostilbene, piceatannol, 6-hydroxyflavone, and kaempferol. Diffracting quality crystals of *Pa*GT2 were obtained only with poor acceptors, resveratrol or pterostilbene, together with UDP-2FGlc. The diffracting quality crystals of *Pa*GT2 with good acceptors were difficult to obtain. Similarly, crystallization of *Pa*GT2 with the donor substrate UDP-glucose also did not yield well diffracting crystals.

The crystal structure of *Pa*GT2 ternary complex with UDP-2FGlc and resveratrol was refined to 2.60 Å resolution (Fig. 2A, Table S1). The asymmetric unit of the ternary complex contained three independent *Pa*GT2 molecules (Fig. S4A). The loop connecting N-and C-terminal domains as well as the loop that caused dimerization of apo-*Pa*GT2 were both disordered in the structure of complex-*Pa*GT2 molecules. The overall structures of all the three molecules were similar with rmsds of 0.46 (chain A and B), 0.56 (chain A and C), and 0.35 Å (chain B and C) for 350, 361, and 358 Cα atoms, respectively. The ternary complex structure of *Pa*GT2 showed shift in N and C-terminal domains towards the substrates. This movement upon binding of substrates is a well-known feature of GT-B fold enzymes (Fig. S5B) (2, 25, 29).

**Fig. 2.**
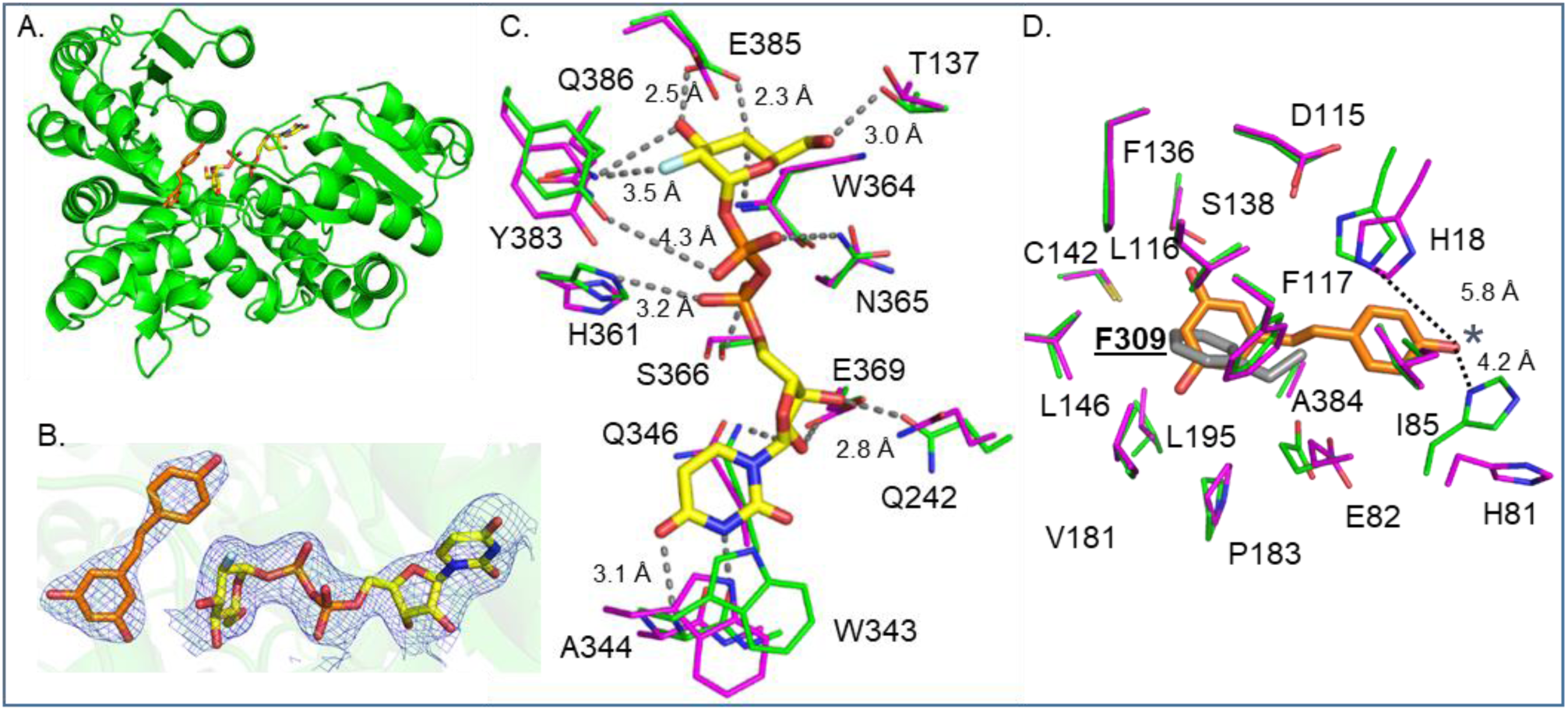
The structure of *Pa*GT2 and its interaction with the substrates. (A) Structure of *Pa*GT2 complexed with resveratrol (orange) and UDP-2FGlc (yellow). (B) Sigma-A-weighted 2*F*o-*F*c electron density maps contoured at 1σ for resveratrol and UDP-2FGlc in chain B of *Pa*GT2 complexed with resveratrol and UDP-2FGlc. (C) Residues involved in binding UDP-2FGlc in complex *Pa*GT2 chain B (green) compared with corresponding residues in apo-*Pa*GT2 (magenta). (D) Residues involved in binding resveratrol in complex *Pa*GT2 chain B (green) compared with corresponding residues in apo-*Pa*GT2 (magenta). The 4’OH group on resveratrol (analogous to glucosylation site on piceatannol) is indicated with blue (*). Phenylalanine (F309) from the opposite molecule occupying the acceptor pocket is in grey color and labelled bold with underline.

The structure of *Pa*GT2 complexed with UDP-2FGlc and pterostilbene was refined to 2.65 Å resolution (Fig. S4B, Table S1). The overall structure of this complex was similar to the structure of *Pa*GT2 complexed with resveratrol and UDP-2FGlc (rmsd 0.80 Å for 1251 Cα atoms).

### Donor binding

UDP-2FGlc was bound in the donor binding pocket on the C-terminal domain of *Pa*GT2. The electron density of UDP-2FGlc was clearly observed in all three molecules in the asymmetric unit of both complex structures (Figs. 2B, S5). UDP-2FGlc in *Pa*GT2 was mainly stabilized by the interaction with residues from the PSPG motif (Figs. 2C, S8C) extending from Trp343 to Gln386 (Fig. S10). The sidechain of Trp343 shifted toward the uracil ring in the UDP-2FGlc bound structure (Fig. 2C). This highly conserved Trp is observed to flip and form a π-stacking interaction with the uracil ring of UDP-sugar donor in other known UGT structures (3, 4). However, in the crystal structure of *Pa*GT2, the indole ring of Trp343 was not flipped after binding the donor (Fig. S7) and no such π-stacking interaction was observed. The O4 oxygen and the N3 nitrogen of uracil ring formed hydrogen bond with the main chain nitrogen and oxygen atoms of Ala344, respectively. The oxygen atoms in the ribose ring interacted with Glu369. The O2 oxygen and the O3 oxygen on the ribose ring formed hydrogen bond with Gln346 and Gln242, respectively. Gln242 in complex-*Pa*GT2 was observed to move towards UDP-2FGlc as compared to apo-*Pa*GT2 (Fig. S7). The interaction between Gln242 and the ribose moiety could be important in *Pa*GT2 for the stabilization of the sugar donor. The O1A and the O2A oxygen atoms on α-phosphate formed hydrogen bond with Ser366 and His361, respectively. The oxygen atoms O1B and O2B on β-phosphate interacted with Asn365 and Tyr383, respectively. Gln386 stabilized the 2F fluorine and the O3 oxygen atoms on the glucose moiety. The O3 and the O4 oxygen atoms of the glucose ring formed hydrogen bonds with Glu385. The O4 oxygen atom of the glucose also formed hydrogen bond with the main chain nitrogen of Trp364. Thr137 and Asn365 stabilized the O6 atom on the glucose moiety. The residues that interacted with the sugar donor are highly conserved among the plant UGTs. The relevance of the residues that interact with the sugar donor molecule has been studied in different UGTs and has been shown to impair UGTs upon mutation (2, 3, 25).

### Acceptor binding

The acceptor binding site in N-terminal domain of *Pa*GT2 was made up mainly of hydrophobic residues. The crystal structures showed the electron densities for both resveratrol and pterostilbene in their respective complexes (Figs. 2B, S5). The electron density for resveratrol was clearer in one of the *Pa*GT2 molecules (chain B) in the asymmetric unit (Fig. 2B). The electron density for resveratrol in other two *Pa*GT2 molecules of the asymmetric unit was observed only for the resorcinol moiety (ring A). The electron density of pterostilbene was clearer as compared to that of resveratrol (Fig. S6B). However, the electron densities of phenol moieties (ring B) of pterostilbene were still weaker than the electron densities of ring A. The electron densities of both resveratrol and pterostilbene clearly indicated that ring A of stilbenoids occupied the inner space of the acceptor binding pocket and ring B pointed towards the solvent.

The acceptor binding pocket in *Pa*GT2 was formed by hydrophobic residues Ile85, Leu116, Phe117, Phe136, Leu146, Val181, Pro183, Leu195, Ala384, and some polar residues His18, His81, Glu82, Cys142, and Ser138 (Figs. 2D, S7A, S7B). Ring A in stilbenoids were stabilized mainly by hydrophobic interactions in the interior of the acceptor pocket and through a CH-π stacking interaction with Leu116. Comparison of apo and stilbenoid-complex *Pa*GT2 structures showed that the acceptor binding site in apo-*Pa*GT2 was occupied by residues from the dimerizing loop. It is noteworthy that the aromatic ring of Phe309 from the opposite molecule was placed at the position occupied by ring A of stilbenoids in complex-*Pa*GT2 (Fig. 2D). Two polar residues Ser138 and Cys142 were at the mean distance of 3.36±0.03 Å and 4.36±0.18 Å, respectively, from the 3OH of resveratrol (SI Table 3). In the structure with pterostilbene, these residues were at the mean distance of 4.06 ± 0.16 Å and 4.29 ± 0.41 Å from the 3-methoxy group of pterostilbene, which was similar to the case of resveratrol despite larger size of pterostilbene. While Cys142 was found only in *Pa*GT2 (Fig. S10), Ser138 is well conserved among the plant UGTs and is closer to Glu385 instead of the acceptors. These observations indicated that the acceptors lacked efficient polar interactions with polar residues of enzyme present at the rear side of the acceptor binding pocket. The highly conserved catalytic histidine present in plant UGTs was recognized as His18 in *Pa*GT2. The sidechain of highly conserved Asp115 was close to His18. The catalytic histidine removes the proton from glycosylation site of acceptor for nucleophilic attack on C1 carbon of carbohydrate moiety on sugar donor. The conserved aspartate at this position is considered to interact with catalytic histidine and assist it in abstraction of proton from the glycosylation site of acceptors (2, 4). In the crystal structures, the mean distance between His18 and the 4’OH groups on resveratrol and pterostilbene, which corresponded to the glycosylation site of piceatannol, were observed to be 5.36 ± 0.23 Å and 6.06 ± 0.03 Å, respectively (SI Table 3). These distances were long for proper hydrogen bonding. Ring B of stilbenoids were also stabilized by hydrophobic interactions, mainly from Ile85 and Ala384. The closest polar residue His81 was at the mean distance of 3.76 ± 0.21 Å and 4.67 ± 0.82 Å from 4’OH of resveratrol and pterostilbene, respectively (SI Table 3). Glu82, positioned next to His81, was also not suitable for formation of hydrogen bond with these stilbenoids in the crystal structures. However, His81 and Glu82 could be important for stabilization of substrates, such as piceatannol and kaempferol. The importance of His81 and Glu82 in catalysis was elucidated by docking simulation for piceatannol and kaempferol in the *Pa*GT2 structure (Fig. 3). As was observed in the crystal structures with stilbenoids, His81 was closer to 4’OH of modeled piceatannol compared to the conserved catalytic residue His18 (SI Table 4). In contrast, His18 was closer to the 3OH glucosylation site in modeled kaempferol than His81, while H81 could interact with 4’OH also in kaempferol. The docking result also suggested that Glu82 could form a hydrogen bond with 3’OH in piceatannol and be close to the O1 oxygen in kaempferol. Indeed, polar residues at corresponding position were reported in other UGTs, such as Gln84 in *Vv*GT1(2) and Glu88 in *Pt*UGT1(26), stabilized the respective acceptor molecules by forming a hydrogen bond. Thus, residues located around the outer surface of the acceptor binding site are thought to be involved in the stabilization of substrates. Contrary, due to the lack of 3’OH group on ring B, resveratrol and pterostilbene would have weaker hydrogen bonding interactions than piceatannol, resulting in the flexible structures of stilbenoids in the acceptor binding pocket. The weak stabilization of these stilbenoids could be the reason for their negligible glucosylation and poor electron density of ring B in the crystallographic data.

**Fig. 3.**
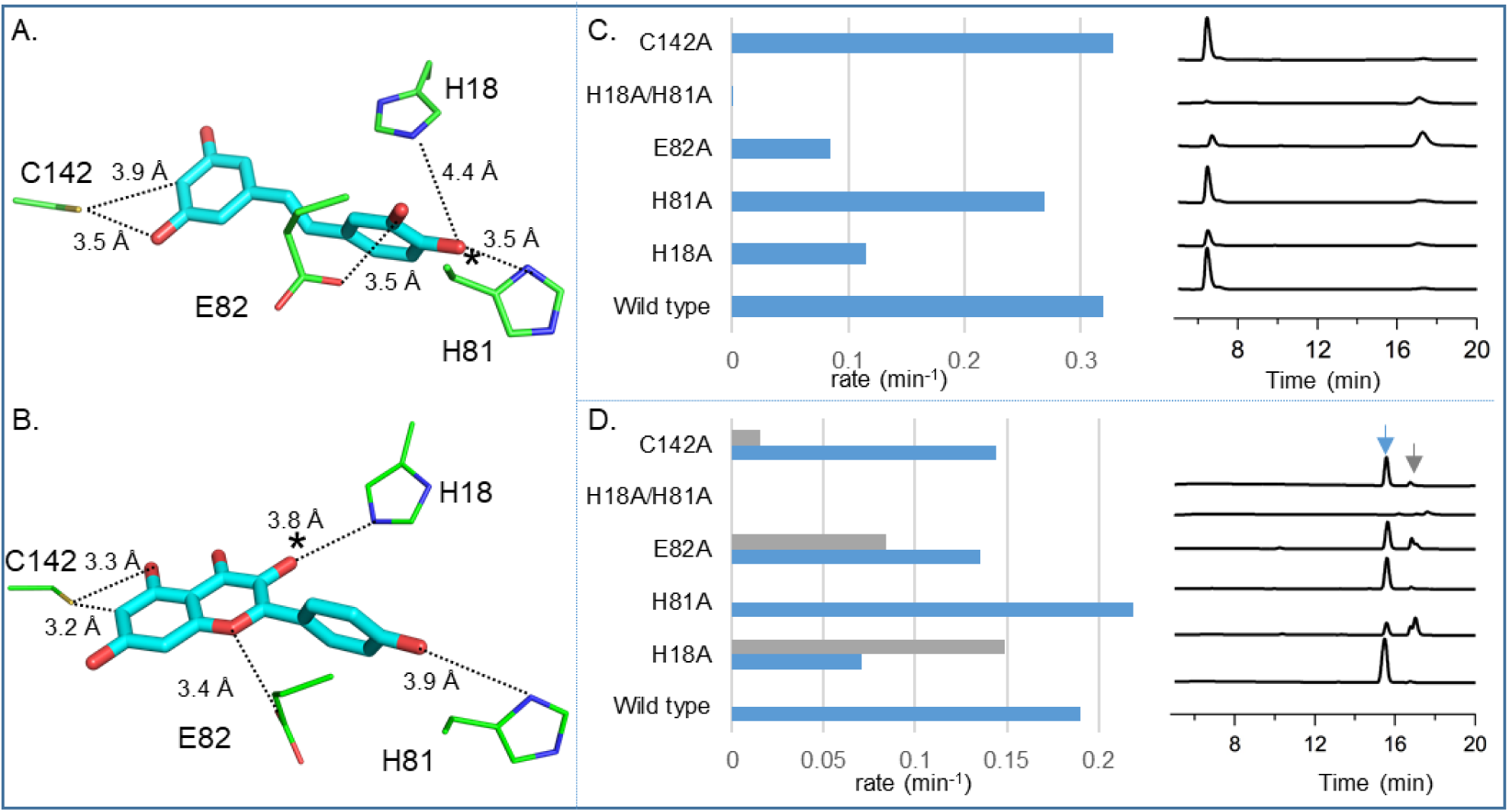
Docking and mutational analysis. (A, B) Conformation of (A) piceatannol and (B) kaempferol in *Pa*GT2 from docking model prepared by software PyRx. Glucosylation sites are indicated with *. (C, D) The result from glucosylation activity and HPLC profile of *Pa*GT2 mutants for (C) piceatannol and (D) kaempferol is shown. For kaempferol, blue bars/arrow indicate the glucosylation product kaempferol-3-*O-*β-glucoside and grey bars/arrow indicate other glucoside products.

### Mutagenesis studies

In order to confirm the involvement of the residues in catalysis and substrate recognition, *Pa*GT2 mutants were prepared and their glucosylation activities were evaluated. The residues that were assumed to directly interact with the acceptors were mutated for the mutational study. The list of *Pa*GT2 mutants and their glucosylation activity data are listed in Table 1. For the functional assay, piceatannol was used as the representative stilbenoid as wild type (WT) *Pa*GT2 had very low activity with other stilbenoids. Functional assays were also conducted with kaempferol to determine the difference between the glucosylation of stilbenoids and flavonoids.

The substitution of Cys142 with a smaller alanine residue increased the *K*_m_ value by 1.7-fold but did not decrease the *k*_cat_ value for the production of piceatannol 4’-*O*-β-glucoside. The *K*_m_ value of the Cys142Ala mutant towards kaempferol was comparable with the WT *Pa*GT2, while the *k*_cat_ value was increased by about 40%. Moreover, the Cys142Ala mutant transformed kaempferol not only to kaempferol 3-*O*-β-glucoside but also a small amount of other glucosides, indicating the importance of Cys142 in regioselective glucosylation of kaempferol. The substitution of Cys142 with a smaller amino acid residue appeared to afford space for ring A of piceatannol/kaempferol. The slight increase in acceptor binding pocket could have allowed other conformations of kaempferol, that were not possible in WT *Pa*GT2, and resulted in production of side products.

We also constructed His81Ala and Glu82Ala mutants to examine the role of these residues in glucosylation. For piceatannol glucosylation, both of these mutants exhibited lower *k*_cat_ and higher *K*_m_ values than the WT *Pa*GT2, which indicated their involvement in the efficient formation of the Michaelis complex. In the case with kaempferol, the *K*_m_ value for kaempferol glucosylation was nearly double with the His81Ala mutant. The *K*_m_ value for kaempferol glucosylation by the Glu82Ala mutant was comparable to the WT enzyme but this mutant lost regioselectivity with slight increase in the total *k*_cat_/*K*_m_ value. These results indicated that both His81 and Glu82 were also involved in the glucosylation of kaempferol.

To somewhat surprise, the His18Ala mutant retained glycosylation activity for both piceatannol and kaempferol although His18 was the conserved catalytic residue. For both piceatannol and kaempferol glucosylation, the *K*_m_ value was not affected significantly by the mutation of His18 to Ala. The *k*_cat_ value for piceatannol glucosylation by the His18Ala mutant was 30% of the WT enzyme. This mutant had low regioselectivity and hence produced a mixture of several kaempferol glucosides; however, the total catalytic efficiency was comparable with WT *Pa*GT2. We also prepared a mutant deleting 31 N-terminal residues (*Pa*GT2Δ31), which lacked the conserved histidine (His18). This mutant was able to glucosylate piceatannol as with the His18Ala mutant. The piceatannol glucosylation activity of Δ31 mutant was low compared to His18Ala mutant (Fig. S9) presumably because the N-terminal deletion could have distorted the structure of enzyme and affected its activity. In order to identify the residue that catalyzed the glucosylation reaction in His18Ala and Δ31 mutants, we generated the His18Ala/His81Ala double mutant *Pa*GT2, taking the proximity of His81 towards 4’OH of acceptors in crystal structures and formation of piceatannol 4’-*O*-β-glucoside into considerations. The enzyme assay showed that the His18Ala/His81Ala mutant lost the ability to generate glucosides with both piceatannol and kaempferol. These results indicated that His81 could be another residue in *Pa*GT2 that could catalyze the glucosylation.

## Discussion

*Pa*GT2, a glycosyltransferase from *P. americana*, can glucosylate structurally different polyphenols: stilbenoids and flavonoids (Figs. 1, S2). For the structure-function relationship of *Pa*GT2 we determined the crystal structures of *Pa*GT2 with and without stilbenoids and UDP-2FGlc. The stilbenoids in *Pa*GT2 structures are placed in the highly hydrophobic acceptor binding pocket with ring B pointing outward of the cavity (Fig. 2D). The part of stilbenoids lying in the interior of acceptor binding pocket is stabilized mainly by van der Waals interactions and CH-π stacking interaction with Leu116. The distance between acceptors and Cys142 in the crystal structure is long for effective hydrogen bonding. However, the mutation of Cys142 to Ala showed an increase in the *K*_m_ for piceatannol glucosylation and loss in regioselectivity of kaempferol glucosylation, indicating its role in substrate binding. The interaction of enzyme with ring B of substrates in the crystal structures is weaker. This is due to lack of polar groups in resveratrol/pterostilbene other than 4’OH on ring B. In the case of piceatannol or kaempferol, acceptors are stabilized by His81 and Glu82. In fact, the docking model of piceatannol in *Pa*GT2 indicates that Glu82 could form a hydrogen bond with 3’OH of piceatannol (Fig. 3A). The mutation of Glu82 to Ala reduces the *k*_cat_ value to around 30% of WT *Pa*GT2 and increases the *K*_m_ value for piceatannol glucosylation, supporting the hypothesis that Glu82 stabilizes ring B of piceatannol. Also, mutation of His81 to Ala severely affected both the *K*_m_ and *k*_cat_ values for piceatannol glucosylation. Thus, the low activity of *Pa*GT2 towards resveratrol and pterostilbene is due to weak stabilization of these substrates in the acceptor binding pocket, whereas the presence of methoxy group on 4’ position on rhapontigenin prevented its glucosylation. The docking model of kaempferol indicates Glu82 and His81 are also important for the stabilization of kaempferol (Fig. 3B). The *K*_m_ value for kaempferol glucosylation is almost double in the His81Ala mutant although the catalytic efficiency is not affected. Glu82 is likely to interact with O1 oxygen in kaempferol and stabilize it in the active site. The Glu82Ala mutant affords a mixture of kaempferol glucosides including the 3-*O*-β-glucoside. This suggests that Glu82 stabilizes kaempferol in a particular single conformation in the active site for high regioselective glucosylation of kaempferol. The loss of interaction between O1 oxygen in kaempferol and Glu82 could have allowed different conformations of kaempferol in the active site resulting into its poor regioselective glucosylation. These observations suggest that the residues at position 81 and 82 can make contact with ring C and B of flavonoids, such as kaempferol, to stabilize them in the active site.

Plant UGTs are usually characterized by the presence of a conserved catalytic pair. In *Pa*GT2 the conserved catalytic pair corresponds to His18 and Asp115. However, the His18Ala mutant still shows 30% activity with piceatannol and comparable activity with kaempferol to produce piceatannol 4’-*O*-β-glucoside and a mixture of kaempferol glucosides, respectively. Mutation of the conserved histidine in *Vv*GT1 (His20) (2), UGT85H2 (His21) (4), and *Pt*UGT1 (His26) (26) are reported to result in complete loss of enzyme activity. Another GT from *P. americana, Pa*GT3, also loses the glycosylation activity when the conserved histidine (His20) is mutated to Ala or Asp (22). Moreover, the *Pa*GT2Δ31 mutant, which lacks His18, can glucosylate piceatannol (Fig. S9). It is reported that in a UGT purified from the root of *G. max* (*Gm*IF7GT) lacks 49 N-terminal residues including the conserved catalytic histidine, but can glycosylate its substrate (30). Noguchi *et al*. assumed the presence of the second catalytic residue in *Gm*IF7GT though it was not identified due to the lack of structural information of the enzyme. Therefore, it can be deduced that *Pa*GT2 possesses another active residue that can catalyze glucosylation of the substrates in the absence of His18. The crystal structures of *Pa*GT2 as well as the docking of piceatannol show that His81, compared to His18, is closer to the glucosylation site (4’OH) in stilbenoids (Figs. 2D, 3A). Although this His81 is not the conserved catalytic residue, it is possible that the residue is involved in glucosylation of the substrates especially in the His18Ala mutant as well as in the Δ31 mutant. The role of His81 in catalysis is confirmed by the His18Ala/His81Ala double mutant PaGT2, which shows no glucosylation activity with both piceatannol and kaempferol.

The enzyme assay of individual His18Ala and His81Ala mutants show that His18 is the main catalytic residue for the production of piceatannol 4’-*O*-β-glucoside and kaempferol-3-*O*-β-glucoside (Fig. 3). The increase in the *K*_m_ value for glucosylation of both acceptors show that His81 is involved more in acceptor binding than catalysis. From the enzyme assay data, although His18 is the main catalytic residue for piceatannol glycosylation, it can be assumed that *Pa*GT2 utilizes both His18 and His81 independently for regioselective catalysis (Table 1). For the regioselective kaempferol glucosylation, His18 is the main catalytic residue as both WT and His81Ala has comparable *k*_cat_/*K*_m_ values and forms only kaempferol-3-*O*-β-glucoside. His81 can be considered as a secondary catalytic residue for kaempferol glucosylation when His18 is absent in the active site or when kaempferol binds in conformations different from that in the WT docking model (Fig. 3B). Thus, His81 could have catalyzed the kaempferol glucosylation to produce a mixture of glucosides including kaempferol-3-*O*-β-glucoside in the His18Ala mutant. These facts suggest that not only His18 but also His81 is the catalytic residue in *Pa*GT2.

The studies on plant UGTs are getting broader for the production of dyes, therapeutics, and cosmetics. Our study provides the insights into the catalytic mechanism on one of the non-canonical polyphenol UGTs. The crystal structures of *Pa*GT2 complexed with resveratrol/pterostilbene shed light on the regioselective glycosylation of therapeutically valuable stilbenoids and flavonoids. Moreover, the identification of His81 as an alternative catalytic residue in *Pa*GT2 could be one example for how plant UGTs have evolved their catalytic strategies to adapt a large variety of substrates appearing in the course of changes in their environment. The involvement of His81 in catalysis is also useful to explain the plasticity of UGTs, namely why some UGTs can glycosylate similar substrates at different positions or produce more than one glycosides with single substrate. Our results will provide a basis for the development of tailored biocatalyst for the efficient glycosylation of polyphenols for therapeutic and cosmetic uses.

## Materials and methods

### Expression, purification, and enzyme assay

The *Pa*GT2 gene was cloned into a pCold vector and expressed in *E. coli* BL21 star (DE3). The recombinant protein was purified by using standard metal affinity chromatography followed by anion exchange chromatography. Size exclusion chromatography (SEC) was employed as the final purification step. The protein was then concentrated in the SEC buffer (20 mM Tris-HCl pH 8.0, 100 mM NaCl, 5mM DTT) to 15-20 mg/ml for crystallization. *Pa*GT2 mutants for enzyme assay were expressed and purified using following same protocols.

UGT activity of *Pa*GT2 and mutants were assayed in 50 mM potassium phosphate buffer pH 7.4 at 37°C as described in (20). The reaction mixtures were analyzed by HPLC both for determination of products and for the determination of catalytic constants.

Detailed protocols of expression, purification, and enzyme assay are given in SI text.

### Crystallization, data collection, and crystal structure determination

Details of crystallization and data collection are given in SI text. The structure of apo *Pa*GT2 was solved by molecular replacement using *A. thaliana* UGT72B1 (PDB: 2VCH) as a search model using Molrep in CCP4 (31). Structure of *Pa*GT2 complexes were solved using the apo-*Pa*GT2 structure as a search model. Model building and refinement were performed using COOT (32) and refmac5 (33) in CCP4. Figures were prepared using PyMOL (34) and LigPlot+ (35).

### Molecular docking

Molecular docking of piceatannol and kaempferol were performed using automatic docking program PyRx Virtual Screening tool (36). The crystal structure of *Pa*GT2 in complex with UDP-2FGlc and resveratrol was used as reference. Default parameters were used for controlling the docking process.

## Acknowledgements

The authors wish to thank Dr. K. Fujimoto, Fuji molecular planning co., ltd. for the synthesis of UDP-2FGlc. The authors thank the beamline staffs for their support during data collection on BL44XU at SPring-8 under proposal Nos. 2017A6745, 2017B6745, 2018A6844, and 2018B6844. This work was partly supported by Grant-in-Aid for Young Scientists (B) 17K17862 for YF, Grant-in-Aid for Scientific Research (B) 18H02004 for TI, and Grant-in-Aid for Scientific Research (C) 17K05933 for SO.

## Author contributions

R.M., Y.F., H.H., T.I., and S.O. designed research; R.M., N.S., and S.O. performed the assay and analyzed the results; Taisuke N. established protocols for protein purification and performed preliminary crystallographic analysis; Toru N. contributed cloning of PaGT2; R.M. purified and crystallized proteins; R.M. and Y.F. collected and analyzed crystallographic data; R.M., Y.F., and S.O. wrote the paper with input from all authors.

## Supplementary information

### SI Text

#### Materials and methods

##### Expression and purification of *Pa*GT2

*Pa*GT2 cDNA and pCold vector were amplified by polymerase chain reaction (PCR) using following primers: pCold forward 5’-TAGGTAATCTCTGCTTAAAAGCACAG-3’, pCold reverse 5’-ACCCTGGAAATAAAGATTCTCC-3’ *Pa*GT2 forward 5’-CTTTATTTCCAGGGTATGGAAATGGAAGCACCACTC-3’ and *Pa*GT2 reverse 5’-AGCAGAGATTACCTAGCTTTTGCATTGGCTCCATTTAG-3’.

Amplified *Pa*GT2 cDNA was cloned into pCold vector using Infusion kit (Takara Bio USA, Inc.) following the kit manufacturer’s protocol. The construct contained N-terminal 6x histidine tag followed by TEV protease recognition site. A single colony of BL21 star (DE3), transformed with pCold *Pa*GT2, was inoculated into 2 ml LB medium supplemented with 100 μg/ml ampicillin and grown at 37 °C for about 8 hours as starter culture. 200 μl of the starter culture was introduced into 200 ml LB medium with same concentration of antibiotics and grown overnight at 37 °C. This culture was used to inoculate 1 L culture in the same medium in the next morning and the cells were continued to grow at 37 °C. When the OD600 was ~ 0.4, the temperature was decreased to 15 °C. Expression was induced with isopropyl-β-D-thiogalactopyranoside (IPTG) at 15 °C and OD_600_ ~ 0.6-0.8. After 24 hours, cells were harvested by centrifugation at 9000 ×g for 10 min at 4 °C. Harvested cells were frozen with liquid nitrogen and stored at −80 °C until its use.

The cell pellet was re-suspended in buffer-A (20 mM Tris-HCl pH 8.5, 100 mM NaCl, 5 mM DTT) including 1 tablet of protease inhibitor cocktail (Roche). Cells were lysed by sonication on ice with the following pulse sequence: 15 sec burst, 15 sec rest for a total burst time of 10 min at a power output of 80. Lysed cells were subjected to centrifugation at 20,000 ×g for 30 min at 4 °C. The obtained supernatant was filtered using a 0.45 μm membrane syringe filter. The filtered supernatant was loaded on to a Ni-NTA column (HisTrap HP 5 ml) equilibrated with buffer-A. The column was washed with 10 column volume (CV) of buffer-A. *Pa*GT2 was eluted with a 0-50% gradient of buffer-A supplemented with 300 mM imidazole. Fractions containing *Pa*GT2 were pooled, mixed with TEV protease and dialyzed overnight against buffer-A to remove histidine tag. Dialyzed sample was diluted 10 times using 20 mM Tris-HCl pH 8.5 and loaded on to a HiTrap Q (5 ml) column equilibrated with 20 mM Tris-HCl pH 8.5. *Pa*GT2 was eluted with a linear gradient ranging from 0 to 1 M NaCl in 20 mM Tris-HCl pH 8.5. Fractions containing *Pa*GT2 were pooled, concentrated using Vivaspin and loaded on to Hiload 16/60 Superdex 200 pg size exclusion chromatography (SEC) column equilibrated with SEC buffer (20 mM Tris-HCl pH 8.0, 100 mM NaCl, 5 mM DTT). Fractions containing *Pa*GT2 were pooled, concentrated, and stored at −80 °C.

##### Site-directed mutagenesis

Site-directed mutagenesis was performed using the whole plasmid pCold-*Pa*GT2 and specific oligonucleotide primers listed in SI Table 2. Briefly, the whole plasmid was linearized with mutagenesis primers by PCR. Amplified PCR products were treated with DpnI (New England Biolabs) and purified using NucleoSpin® Gel and PCR Clean-up (MACHEREY-NAGEL), following the manufacturer’s protocol. Purified PCR product were ligated using T4 polynucleotide kinase (Toyobo co.) and ligation high ver. 2.0 (Toyobo co.), and transformed into *E. coli* DH5α. The desired mutation of *Pa*GT2 was confirmed by DNA sequencing.

##### Enzyme assay

WT *Pa*GT2 and all mutants for enzyme assay were expressed and purified as mentioned above. The glucosylation reactions were performed at 37°C in a 200 µl total reaction volume containing 50 mM potassium phosphate buffer (pH 7.4), 50 µM acceptor substrates, 100 µM UDP-glucose, and 5 µM enzyme was incubated at 37°C for 10 minutes. The reaction was stopped by adding 1.5% trifluoroacetic acid, centrifuged at 12,000 rpm for 1 minute and filtered using dismic-13HP. HPLC analysis of the reaction mixtures was performed on Imtact US-C18 (2.0 × 150 mm) reverse-phase column at a flow rate of 0.2 mL/min. Piceatannol and resveratrol glucosylation mixtures were analyzed by isocratic elution, starting with 15% acetonitrile and 85% water for 20 minutes followed by 100% acetonitrile for 10 minutes. Rhapontigenin glucosylation mixtures were also analyzed by isocratic elution, starting with 50% acetonitrile and 50% water for 30 minutes followed by 100% acetonitrile for 10 minutes. Kaempferol glucosylation products were eluted by a linear gradient of acetonitrile starting from 10% acetonitrile and 90 % water to 30% acetonitrile and 70 % water in 20 minutes followed by 100% acetonitrile for 10 minutes. Pterostilbene glucosylation products were also eluted using a linear gradient of 50% acetonitrile and 50% water to 100% acetonitrile in 30 minutes followed by 100% acetonitrile for 10 minutes. The reaction products were identified by comparing retention times of peaks with those of authentic glucosides. To quantitate glucoside products in the reaction mixture, standard curves were generated. For enzyme kinetic studies, acceptor concentrations were varied (25-150 µM). The hyperbolic dependence of glucoside production rates on acceptor concentrations was fitted using the Michaelis–Menten equation to determine *k*_cat_ and *K*_m_.

##### Protein crystallization

Crystallization screening of *Pa*GT2 with and without resveratrol and UDP-2FGlc were performed by the sitting drop vapor diffusion method from 100 nl/100 nl mixture of protein stock (15 mg/ml in 20 mM Tris-HCl pH 8.0, 100 mM NaCl, 5 mM DTT) and well solution. Apo *Pa*GT2 was crystallized using well solution containing 0.1 M magnesium formate, 0.1 M MOPS pH 7.0, and 17% w/v PEG 3350 at 20 °C. Optimal crystals for diffraction were achieved by micro-seeding and hanging drop vapor diffusion from 1 μl/1 μl protein to reservoir solution. Crystals were harvested in same reservoir solution supplemented with 15% ethylene glycol and flash cooled in liquid nitrogen.

Complexes of *Pa*GT2 with UDP-2FGlc and resveratrol/pterostilbene was co-crystallized in the presence of 5 mM UDP-2FGlc and 2 mM resveratrol/pterostilbene in ethanol, mixed 1:1 with a well solution containing 0.11 M potassium citrate, 0.06 M lithium citrate, 0.11 M sodium phosphate, and 23-25% w/v PEG 6000 by hanging drop vapor diffusion. Crystals were harvested in same reservoir solution supplemented with 15% xylitol and flash cooled in liquid nitrogen.

##### Data collection and crystal structure determination

Diffraction data were collected on beamline BL44XU at SPring8 with an MX300HE CCD detector (Rayonix, LLC) and an EIGER X 16M detector (Dectris). Data for apo-*Pa*GT2 was collected to 2.30 Å. *Pa*GT2 complexed with UDP-2FGlc acceptors resveratrol and piceatannol were collected to 2.60 Å and 2.65 Å, respectively. X-ray diffraction data were indexed and scaled using HKL2000 (1) or XDS (2). Structure of apo *Pa*GT2 was solved by molecular replacement using *A. thaliana* UGT72B1 (PDB: 2VCH) as a search model using Molrep in CCP4. Structure of *Pa*GT2 complexes were solved using the solved apo-*Pa*GT2 as a search model. Model building, and refinement were performed using COOT and refmac5, as mentioned previously.

---

**SI Table 1:** Data collection and refinement.

|  | *Pa*GT2 (apo) | *Pa*GT2 + resveratrol +<br>UDP-2FGlc | *Pa*GT2 +<br>pterostilbene +<br>UDP-2FGlc |
| --- | --- | --- | --- |
| **Data collection** |  |  |  |
| X-ray source | SPring-8<br>BL44XU | SPring-8<br>BL44XU | SPring-8<br>BL44XU |
| Detector | Rayonix MX300HE | Rayonix MX300HE | EIGER X 16M |
| Wavelength (Å) | 0.9 | 0.9 | 0.9 |
| Space group | $P2_12_12_1$ | $P2_12_12_1$ | $P2_12_12_1$ |
| Cell dimensions<br>*a*, *b*, *c* (Å)<br>*α*, *β*, *γ* (°) | 91.6 94.8 115.2<br>90.0 90.0 90.0 | 57.3 137.6 208.9<br>90.0 90.0 90.0 | 56.7 136.8 205.1<br>90.0 90.0 90.0 |
| Resolution (Å) | 50.00-2.30 (2.34-2.30) * | 50.00-2.60 (2.64-2.60) | 50.00-2.65 (2.74-2.65) |
| Total reflections | 335208 | 349853 | 311499 |
| Unique reflections | 45171 (2216) | 51665 (2535) | 47340 (4530) |
| $R_{\text{merge}}^{\dagger}$ (%) | 7.2 (95.9) | 8.6 (>100) | 7.1 (>100) |
| $R_{\text{meas}}^{\ddagger}$ (%) | 7.8 (103.0) | 4.0 (51.4) | 7.7 (>100) |
| $I/\sigma(I)$ | 28.5 (2.2) | 25.9 (1.5) | 15.7 (2.0) |
| CC<sub>1/2</sub><sup>§</sup> | (0.757) | (0.689) | (0.694) |
| Completeness (%) | 99.9 (100.0) | 99.0 (100.0) | 99.8 (98.6) |
| Redundancy | 7.4 (7.5) | 6.8 (7.2) | 6.6 (6.7) |
| **Refinement** |  |  |  |
| Resolution (Å) | 36.64-2.30 (2.36-2.30) | 44.85-2.60 (2.66-2.59) | 49.68-2.65 (2.71-2.65) |
| $R_{\text{work}}/R_{\text{free}}$ (%) | 21.99/25.11<br>(30.5/35.0) | 21.57/25.13 (33.4/36.9) | 23.77/25.80<br>(37.7/35.0) |
| RMSD bond length (Å) | 0.0058 | 0.0129 | 0.0139 |
| RMSD bond angles (°) | 1.0341 | 1.3768 | 1.3926 |
| Ramachandran plot (%) |  |  |  |
| Favored | 98.44 | 98.53 | 98.37 |
| Allowed | 1.56 | 1.47 | 1.63 |
| Outliers | 0.00 | 0.00 | 0.00 |
| Average *B*-factor (Å<sup>2</sup>) |  |  |  |
| Protein |  | 79.32 | 86.55 |
| Water | 51.27 |  |  |
| UDP-2FGlc | 47.71 | 78.63 | 84.72 |
| Resveratrol/Pterostilbene |  | 128.08 | 116.43 |
| Molprobity score | 1.30 | 1.44 | 1.49 |
| Molprobity clash score<br>(percentile) | 4.81 (99<sup>th</sup>) | 8.14 (99<sup>th</sup>) | 4.63 (99<sup>th</sup>) |
| PDB ID | 6JEL | 6JEM | 6JEN |

Note: \* Values in parentheses are for the highest resolution shell.
$\dagger R_{\text{merge}} = \sum_{hkl} \sum_i |I_i(hkl) - \langle I(hkl) \rangle| / \sum_{hkl} \sum_i I_i(hkl)$
$\ddagger R_{\text{meas}} = \sum_{hkl} \{N(hkl) / [N(hkl)-1]\}^{1/2} \times | \sum_{hkl} \sum_i |I_i(hkl) - \langle I(hkl) \rangle| / \sum_{hkl} \sum_i I_i(hkl)$ , where $I_i(hkl)$ is the intensity of the $i^{\text{th}}$ observation of reflection $(hkl)$ and $N$ is the redundancy.
$\S CC_{1/2} = (\sigma_y^2 - 1/2 \sigma_\epsilon^2) / (\sigma_y^2 + 1/2 \sigma_\epsilon^2)$ , where, $\sigma_y^2$ the variance of the average intensities across the unique reflections of a resolution shell and, $\sigma_\epsilon^2$ the average of all sample variances of the averaged (merged) intensities across all unique reflections of a resolution shell.

**SI Table 2:**
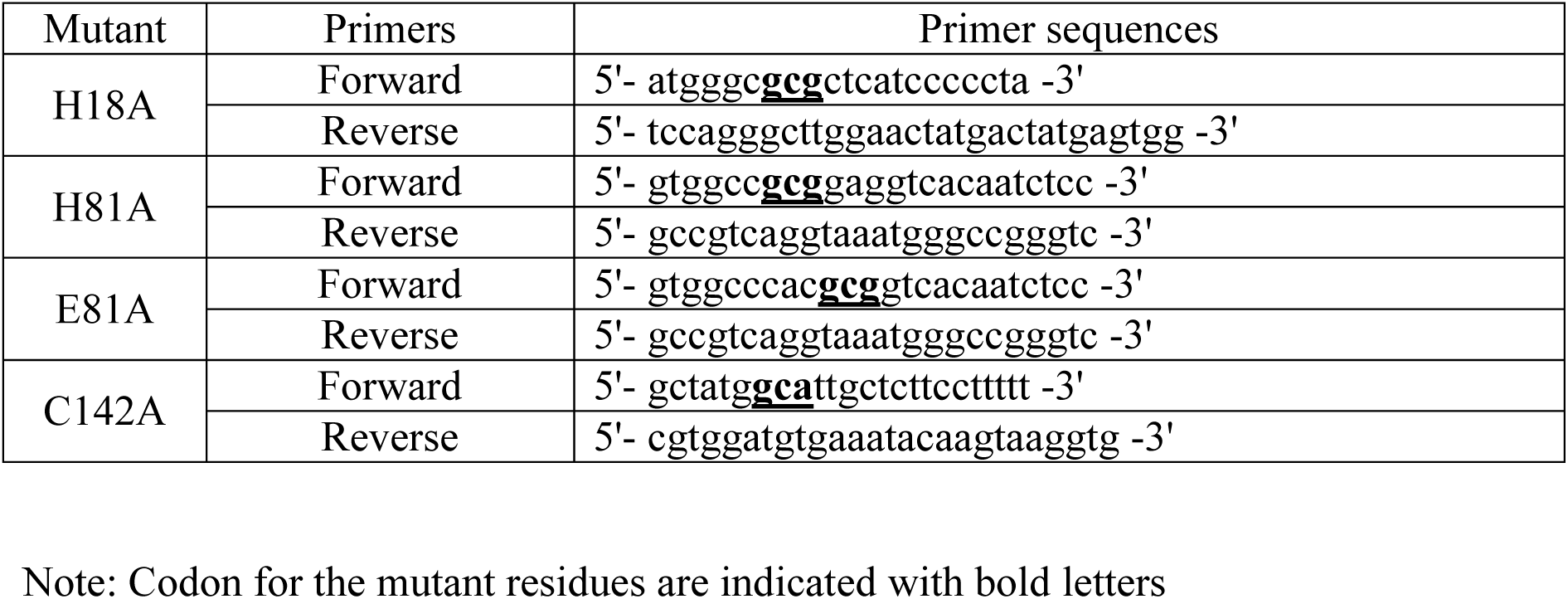
Forward primers used for mutagenesis of *P a*GT2.

**SI Table 3:**
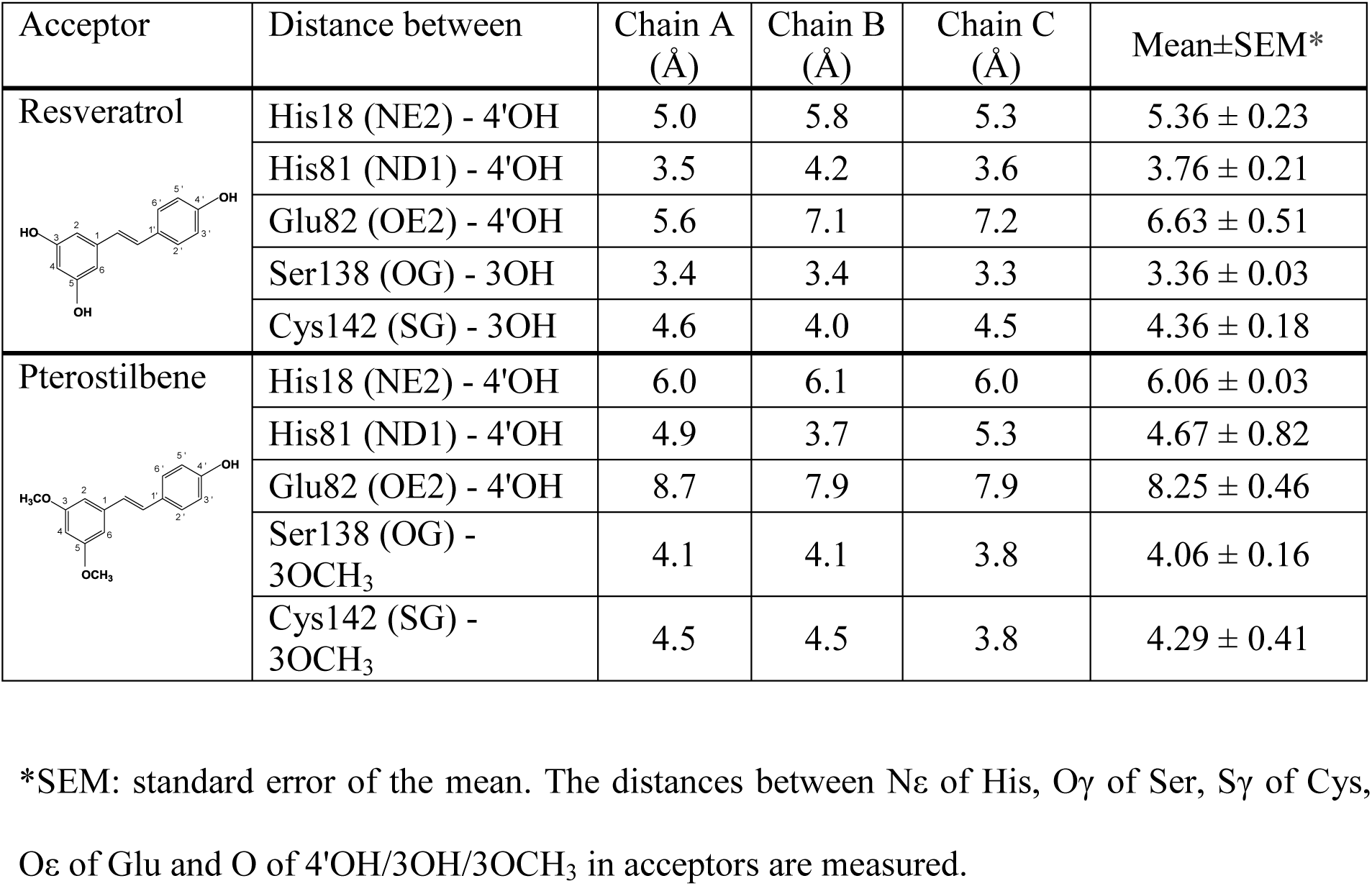
Distance between acceptors and polar residues of enzyme in designated chain in the acceptor binding site of *Pa*GT2.

| Acceptor | Distance between | Chain A<br>(Å) | Chain B<br>(Å) | Chain C<br>(Å) | Mean±SEM* |
| --- | --- | --- | --- | --- | --- |
| **Resveratrol** | His18 (NE2) - 4'OH | 5.0 | 5.8 | 5.3 | 5.36 ± 0.23 |
|  | His81 (ND1) - 4'OH | 3.5 | 4.2 | 3.6 | 3.76 ± 0.21 |
|  | Glu82 (OE2) - 4'OH | 5.6 | 7.1 | 7.2 | 6.63 ± 0.51 |
|  | Ser138 (OG) - 3OH | 3.4 | 3.4 | 3.3 | 3.36 ± 0.03 |
|  | Cys142 (SG) - 3OH | 4.6 | 4.0 | 4.5 | 4.36 ± 0.18 |
| **Pterostilbene** | His18 (NE2) - 4'OH | 6.0 | 6.1 | 6.0 | 6.06 ± 0.03 |
|  | His81 (ND1) - 4'OH | 4.9 | 3.7 | 5.3 | 4.67 ± 0.82 |
|  | Glu82 (OE2) - 4'OH | 8.7 | 7.9 | 7.9 | 8.25 ± 0.46 |
|  | Ser138 (OG) -<br>3OCH<sub>3</sub> | 4.1 | 4.1 | 3.8 | 4.06 ± 0.16 |
|  | Cys142 (SG) -<br>3OCH<sub>3</sub> | 4.5 | 4.5 | 3.8 | 4.29 ± 0.41 |

\*SEM: standard error of the mean. The distances between Nε of His, Oγ of Ser, Sγ of Cys, Oε of Glu and O of 4'OH/3OH/3OCH<sub>3</sub> in acceptors are measured.

**SI Table 4:**
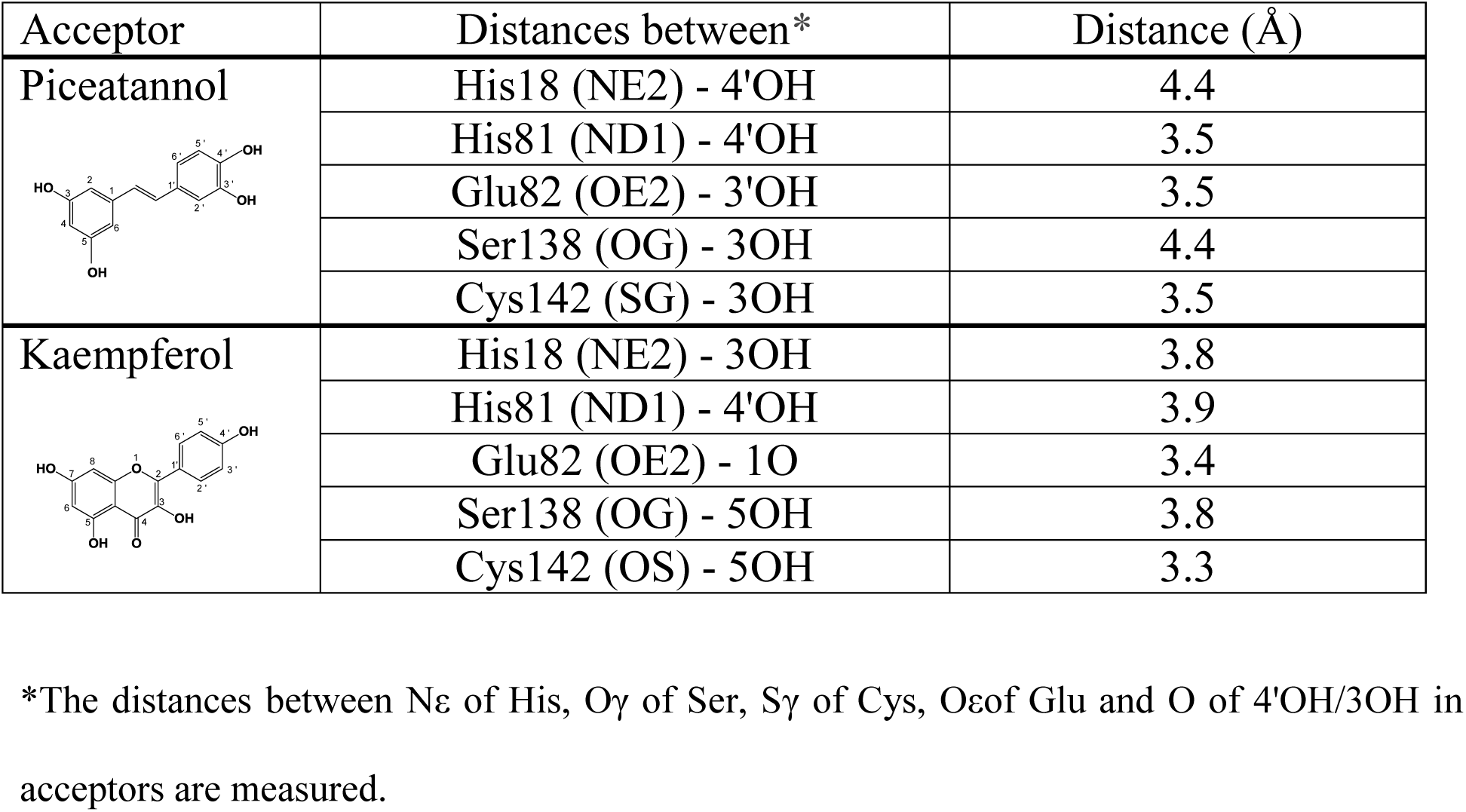
Distance between acceptors and polar residues of *Pa*GT2 in the docking model.

| Acceptor | Distances between* | Distance (Å) |
| --- | --- | --- |
| **Piceatannol** | His18 (NE2) - 4'OH | 4.4 |
|  | His81 (ND1) - 4'OH | 3.5 |
|  | Glu82 (OE2) - 3'OH | 3.5 |
|  | Ser138 (OG) - 3OH | 4.4 |
|  | Cys142 (SG) - 3OH | 3.5 |
| **Kaempferol** | His18 (NE2) - 3OH | 3.8 |
|  | His81 (ND1) - 4'OH | 3.9 |
|  | Glu82 (OE2) - 1O | 3.4 |
|  | Ser138 (OG) - 5OH | 3.8 |
|  | Cys142 (OS) - 5OH | 3.3 |

\*The distances between Nε of His, Oγ of Ser, Sγ of Cys, Oεof Glu and O of 4'OH/3OH in acceptors are measured.

**S1.**
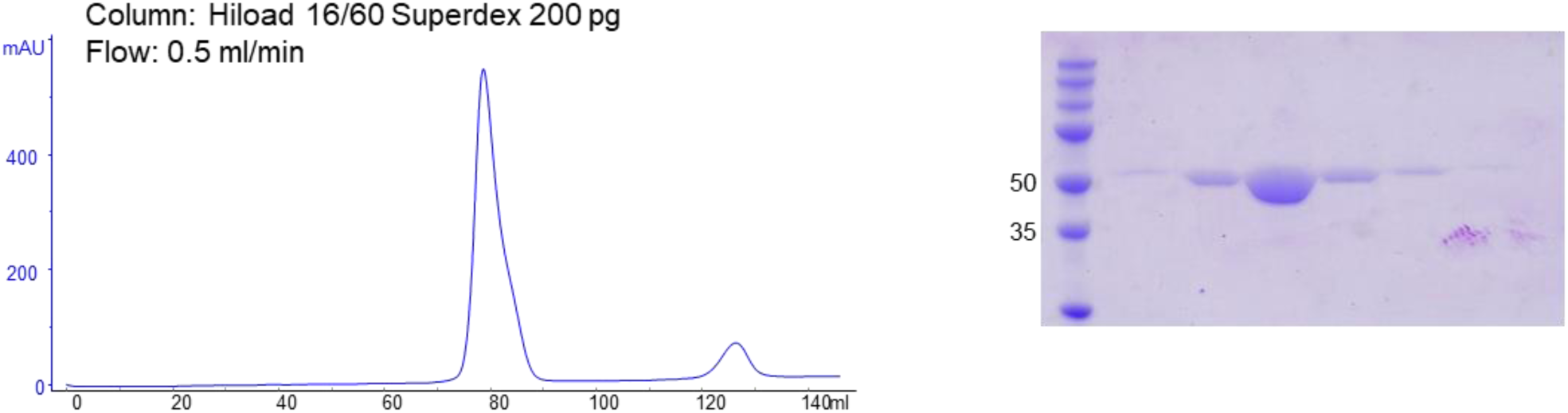
*Pa*GT2 is monomer in solution. (A) Size exclusion chromatography (SEC) profile of *Pa*GT2. The single peak obtained in the chromatogram suggests that the enzyme is monomer in solution. (B) SDS-PAGE of *Pa*GT2 after SEC. The theoretical molecular mass of purified *Pa*GT2 is 51.3 kDa.

**S2.**
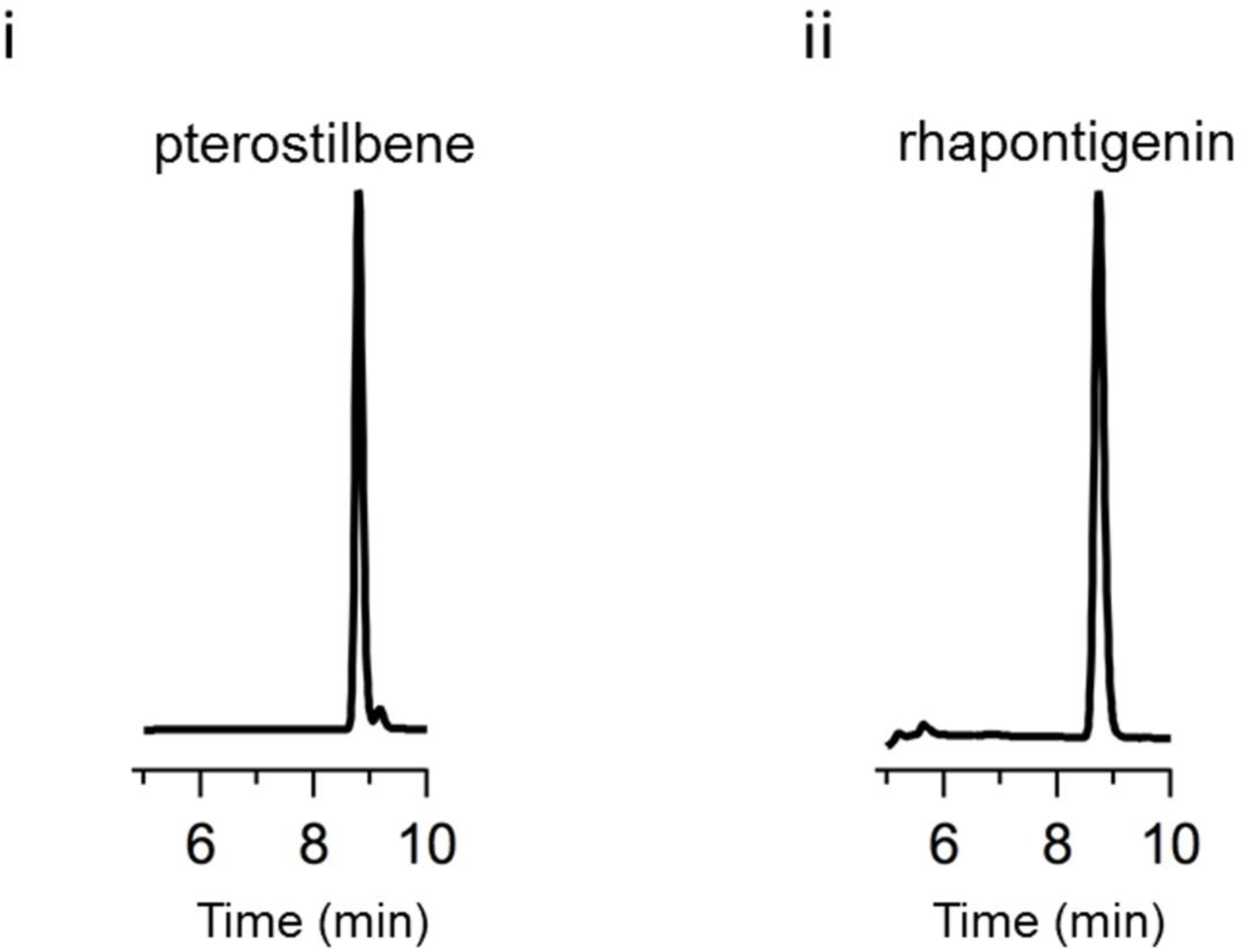
HPLC analysis of aglycones glucosylation by *Pa*GT2. HPLC analysis of i. pterostilbene and ii. rhapontigenin indicated that these are not substrates for glucosylation by *Pa*GT2.

**S3.**
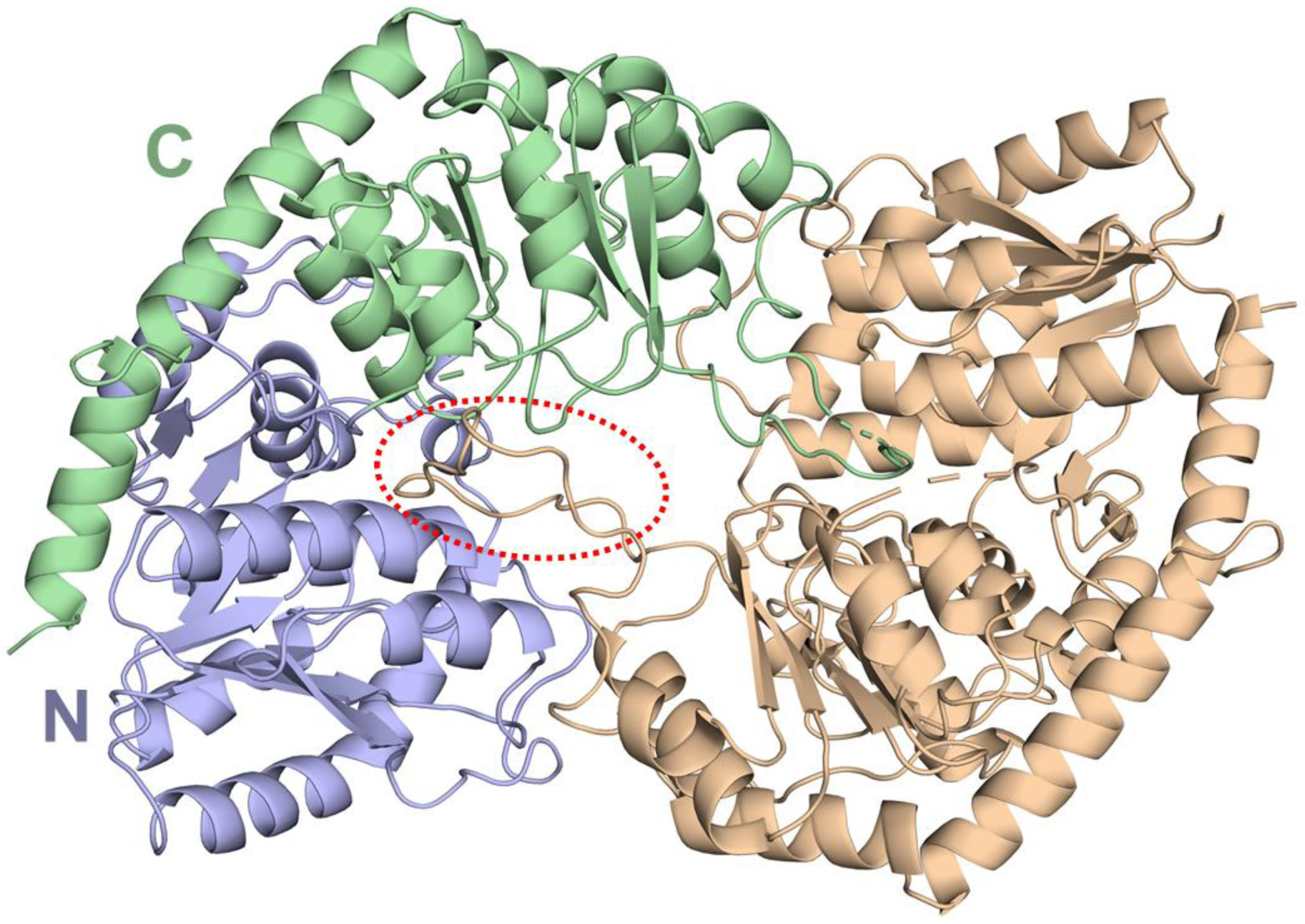
Overall structure of apo-*Pa*GT2. There are two *Pa*GT2 molecules in the asymmetric unit dimerized by insertion of the loop (marked with red oval). The N-terminal (light-blue) and C-terminal (green) domains are indicated in one of the molecule.

**S4.**
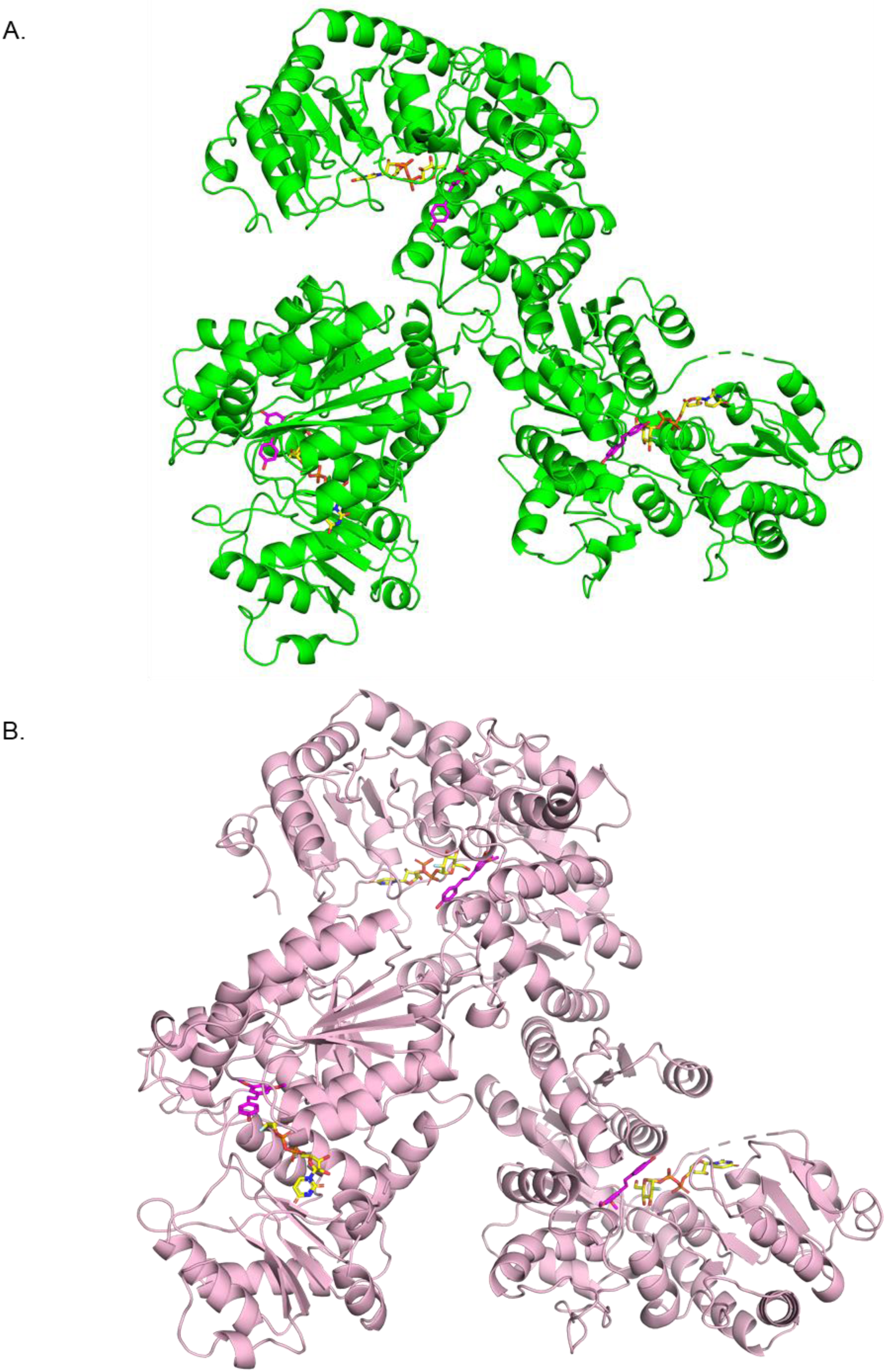
Structure of *Pa*GT2 complexed with substrates. Overall structure of *Pa*GT2 with UDP-2FGlc and (A) resveratrol and (B) pterostilbene in the asymmetric unit.

**S5.**
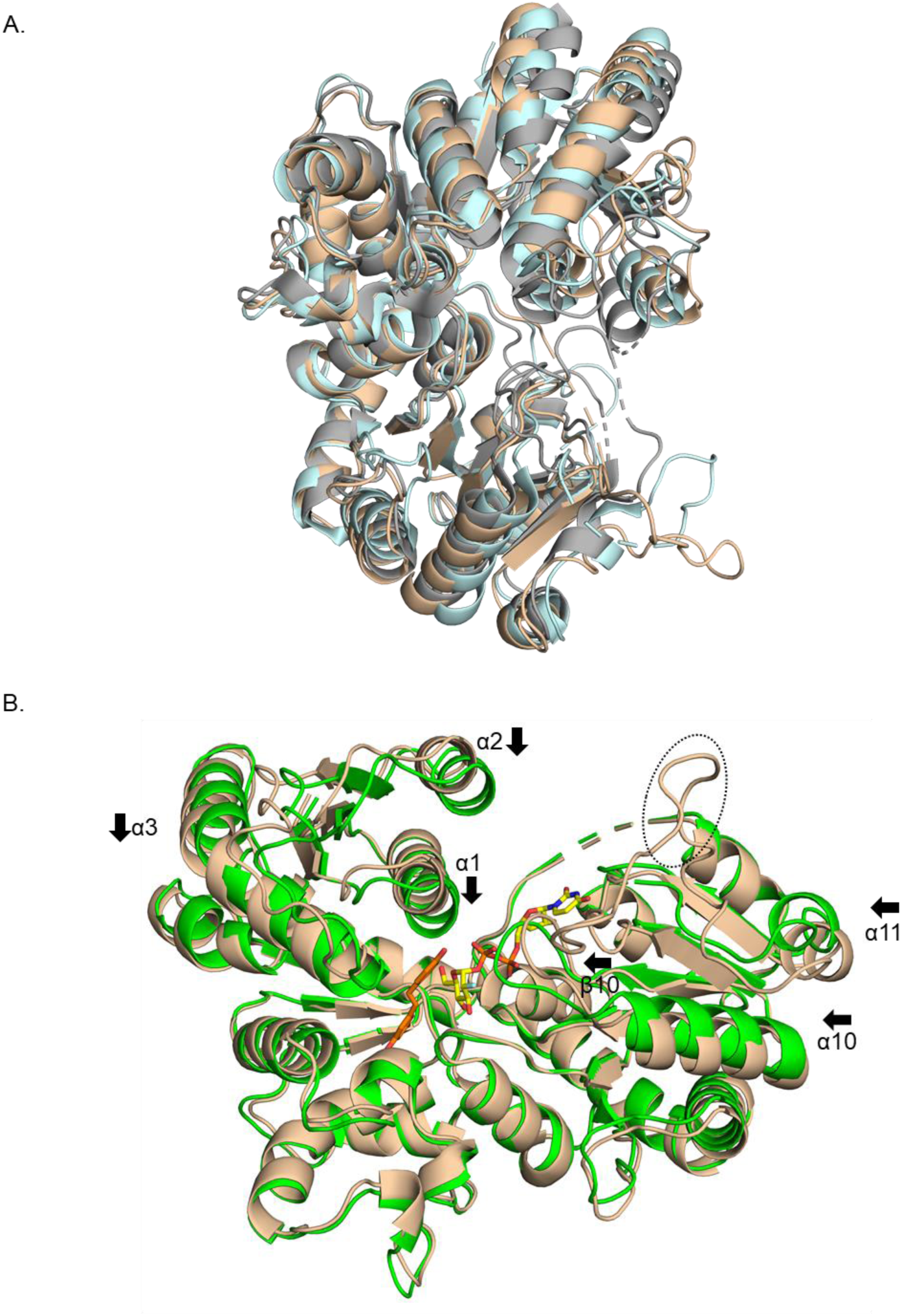
Comparison between other UGT, and *Pa*GT2 structures. (A) Comparison of *Pa*GT2 (light brown), UGT72B1 (grey) and *Pt*UGT1 (light-cyan). (B) Comparison of apo (light brown) and *Pa*GT2 (green) with resveratrol (orange) and UDP-2FGlc (yellow). The loop that causes dimerization of apo-*Pa*GT2 in crystal structure is indicated in black oval. The distinctly shifted secondary structures after binding of substrates are labelled and indicated by arrow heads.

**S6.**
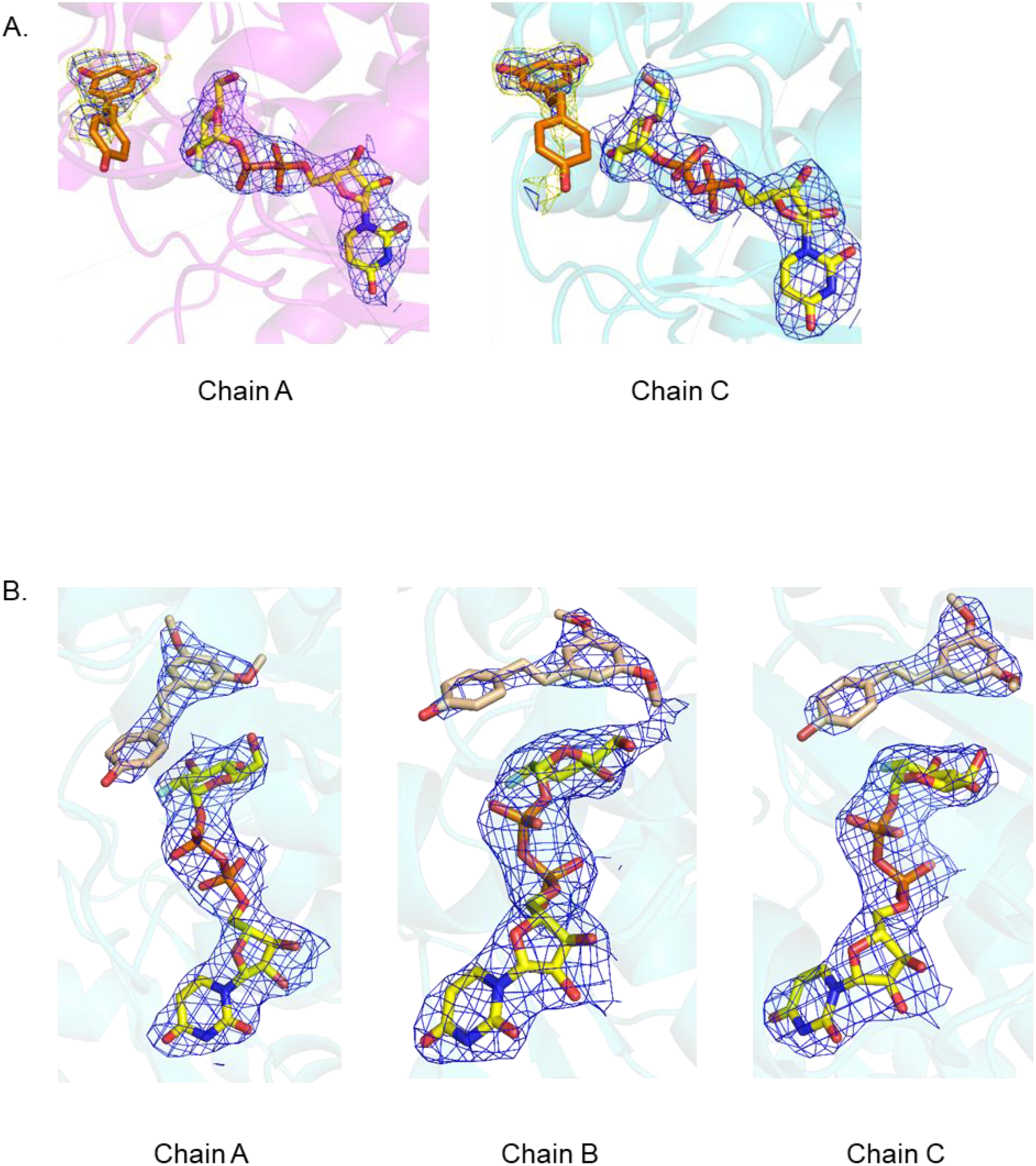
Observed electron densities of substrates. (A) Sigma-A-weighted 2*F*o-*F*c electron density maps contoured at 1σ (blue) and 0.7σ (yellow) for resveratrol and UDP-2FGlc. (B) Sigma-A-weighted 2*F*o-*F*c electron density maps contoured at 1σ for pterostilbene and UDP-2FGlc.

**S7.**
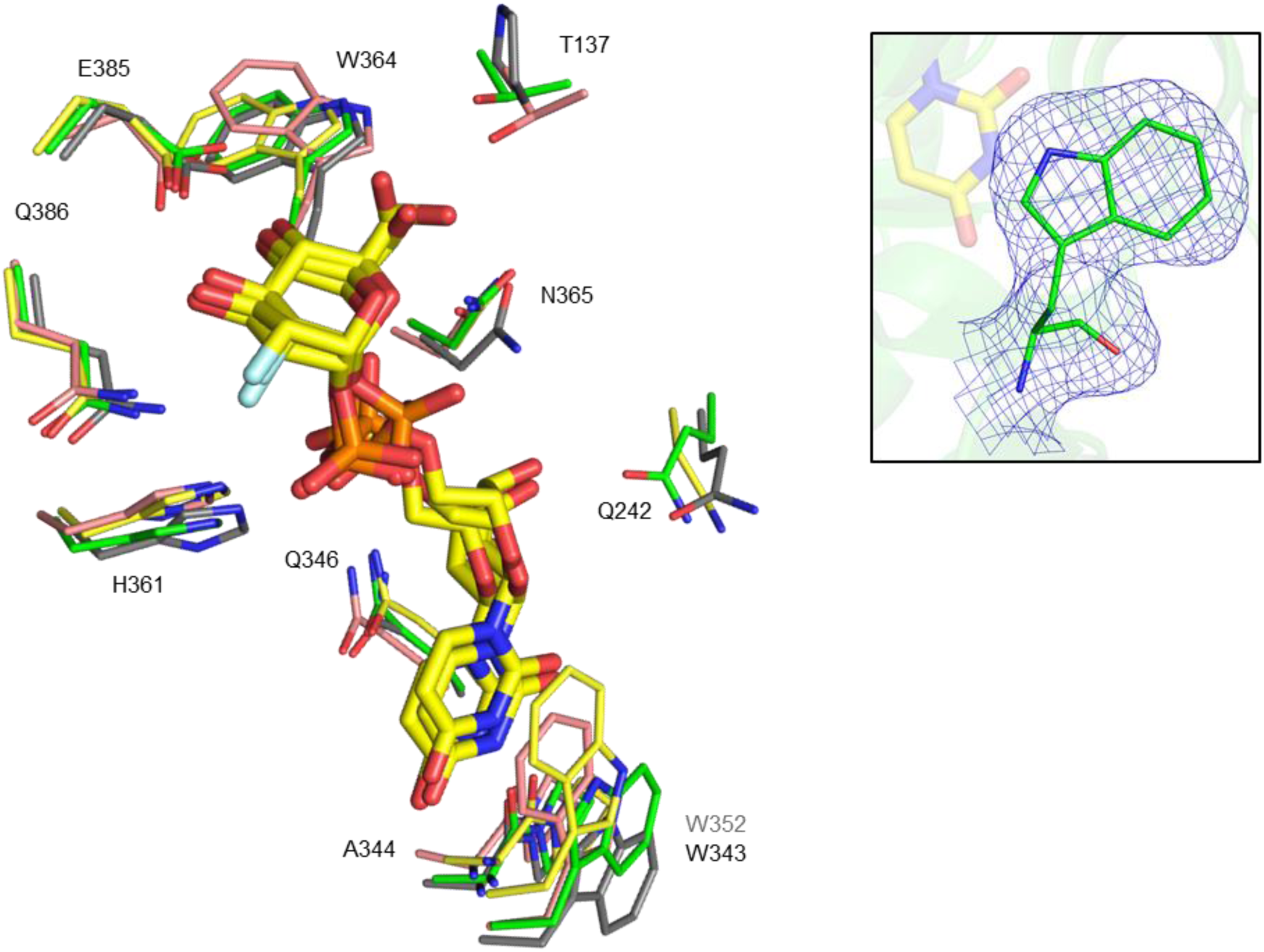
Comparison of donor binding in *Pa*GT2 with other UGTs. Comparison of residues binding UDP-2FGlc (yellow) in *Pa*GT2 (green), *Vv*GT1 (pale red), *Pt*UGT1 (grey) and UGT71G1 (yellow) shows the UDP-glucose binding residues are highly conserved. Residue labels are corresponding to *Pa*GT2. Indole ring of W343 is flipped and sidechain of Q242 is near to UDP-2FGlc in *Pa*GT2. The crystal structure of *Pt*UGT1 (PDB ID: 5NLM) does not contain donor substrate, whose W352 (grey) has same orientation as the W343 of *Pa*GT2. Sigma-A-weighted 2*F*o-*F*c electron density map of W343 contoured at 1.0σ (blue mesh) is shown in inset.

**S8.**
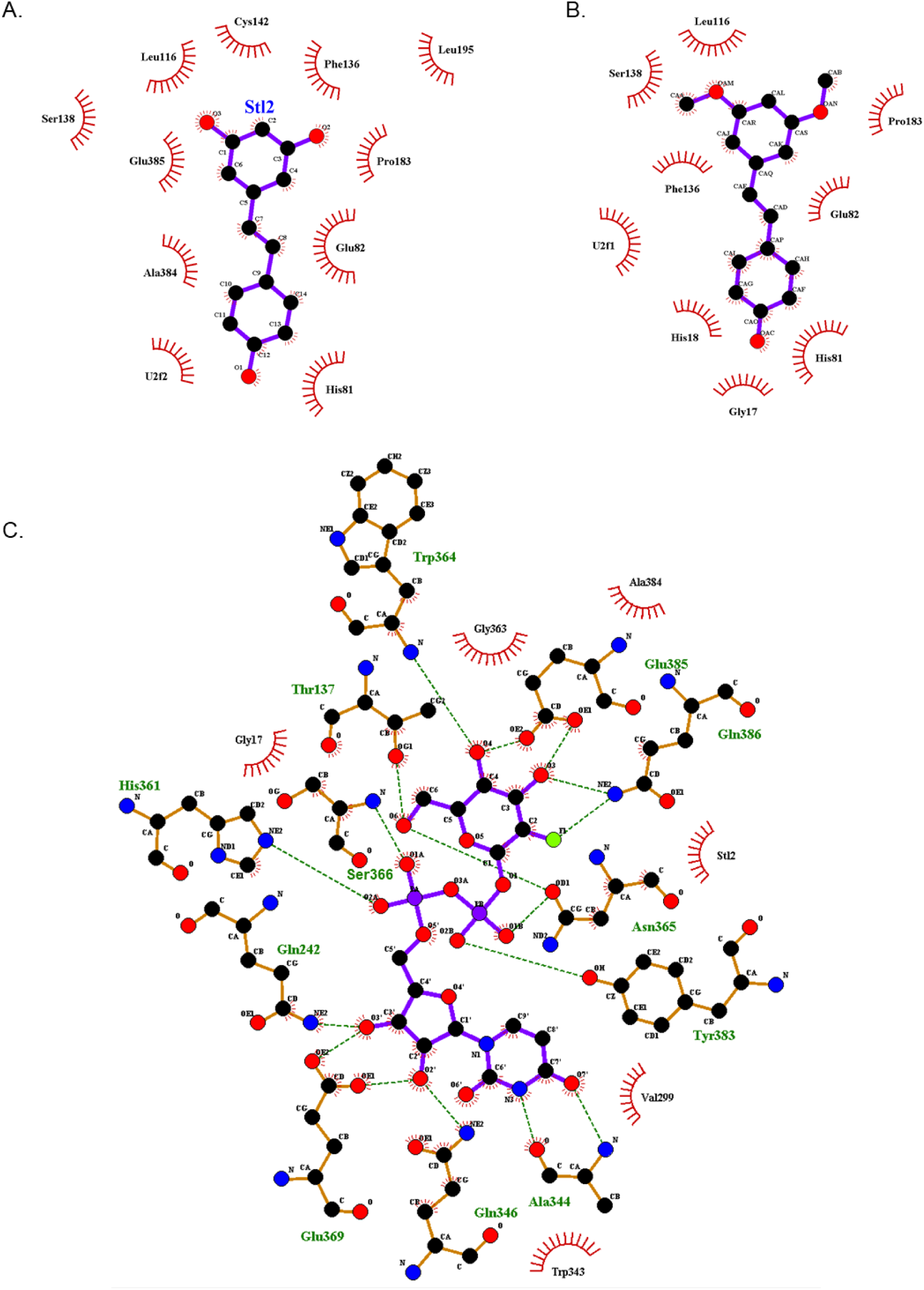
Interaction of *Pa*GT2 with substrates. Interaction of *Pa*GT2 with (A) resveratrol, (B) pterostilbene and (C) UDP-2FGlc. The stilbene acceptors are stabilized mainly by hydrophobic interactions. These figures were drawn using LigPlot^+^.

**S9.**
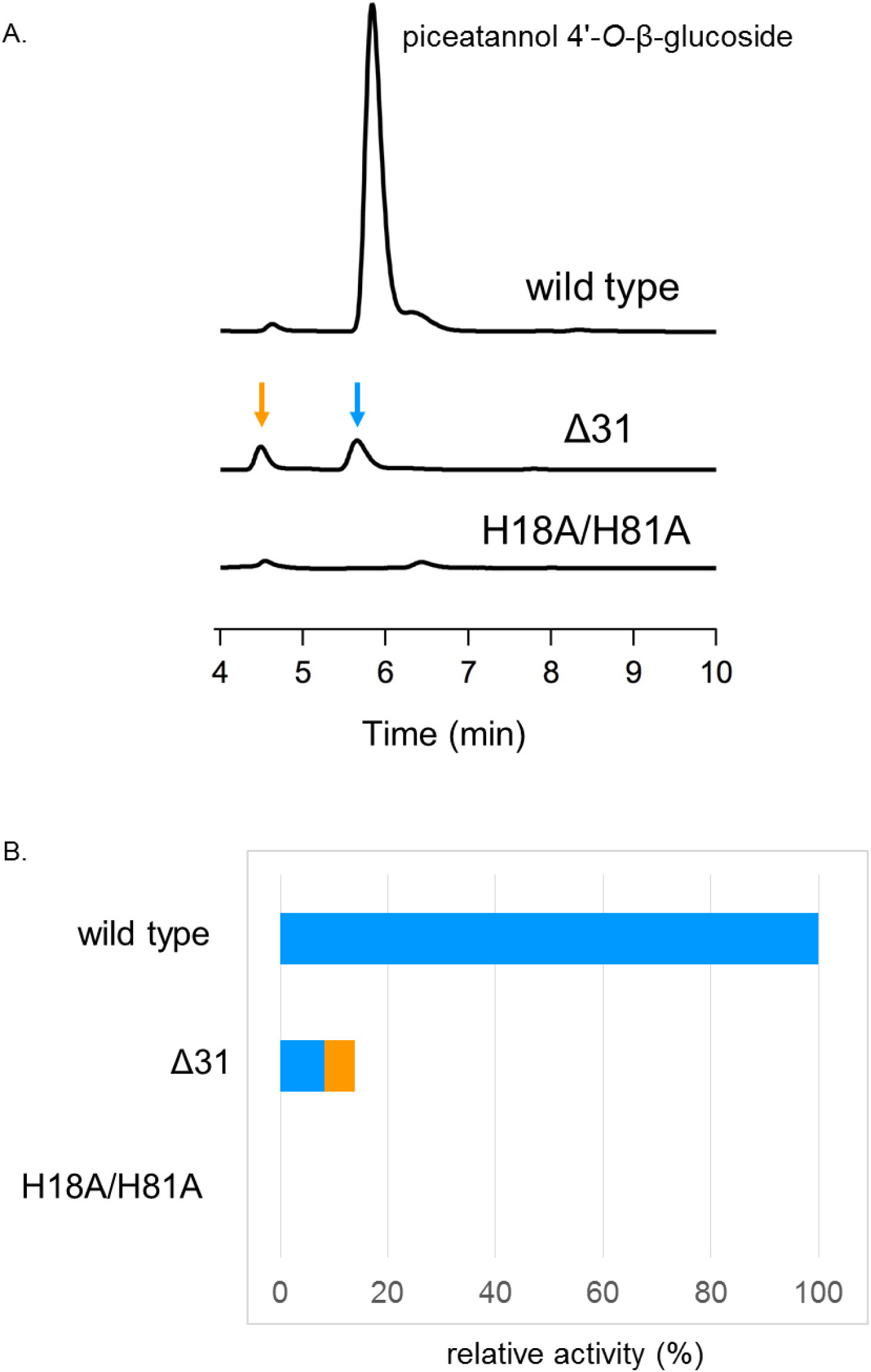
Glycosylation activity of *Pa*GT2 Δ31. (A) HPLC profiles of piceatannol glucosylation. (B) Comparison of the piceatannol glucosylation activity. Relative activity was calculated based on the production of piceatnnnol 4’-*O*-glucoside (blue) and the side product (orange) shown by arrows in (A).

**S10.**
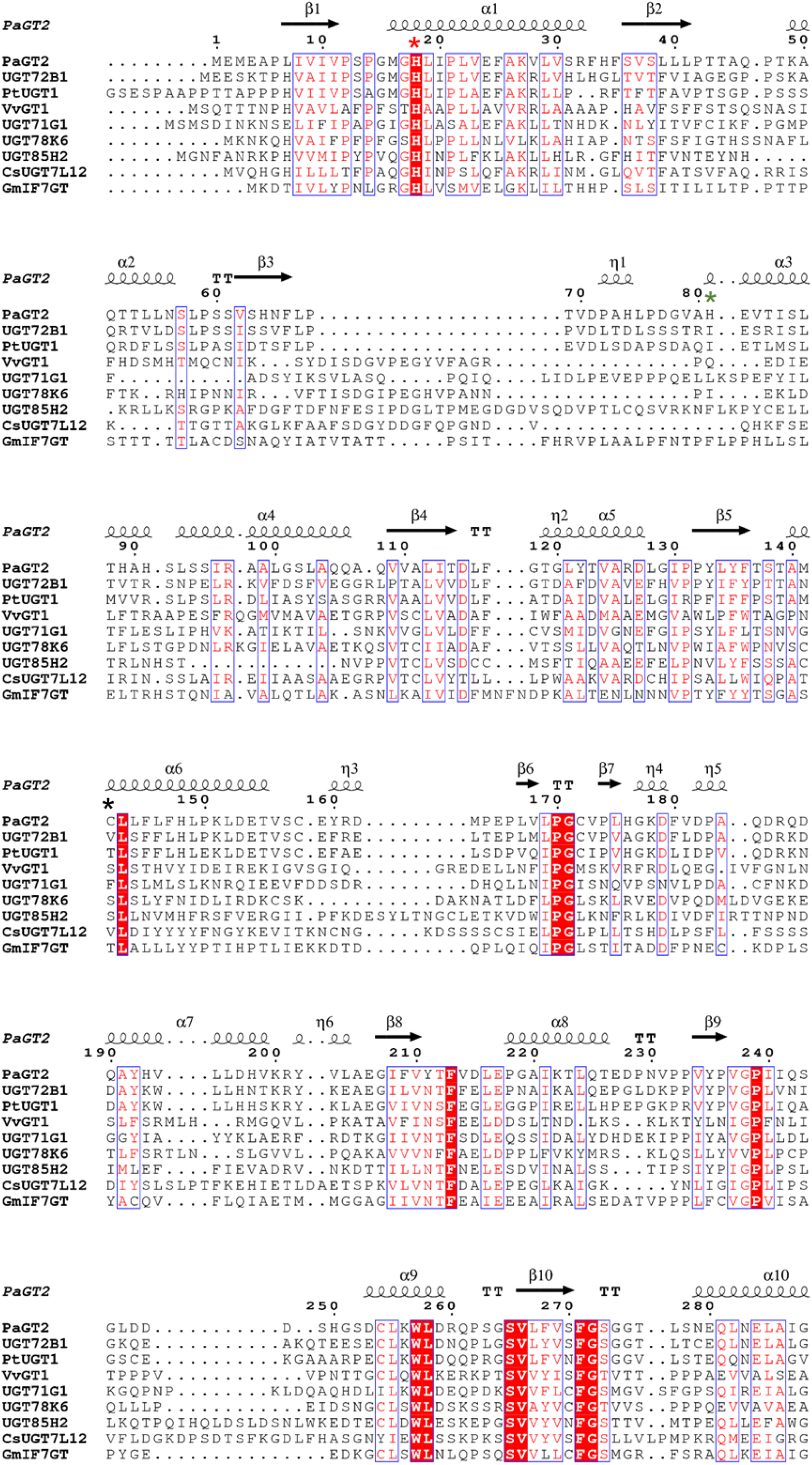
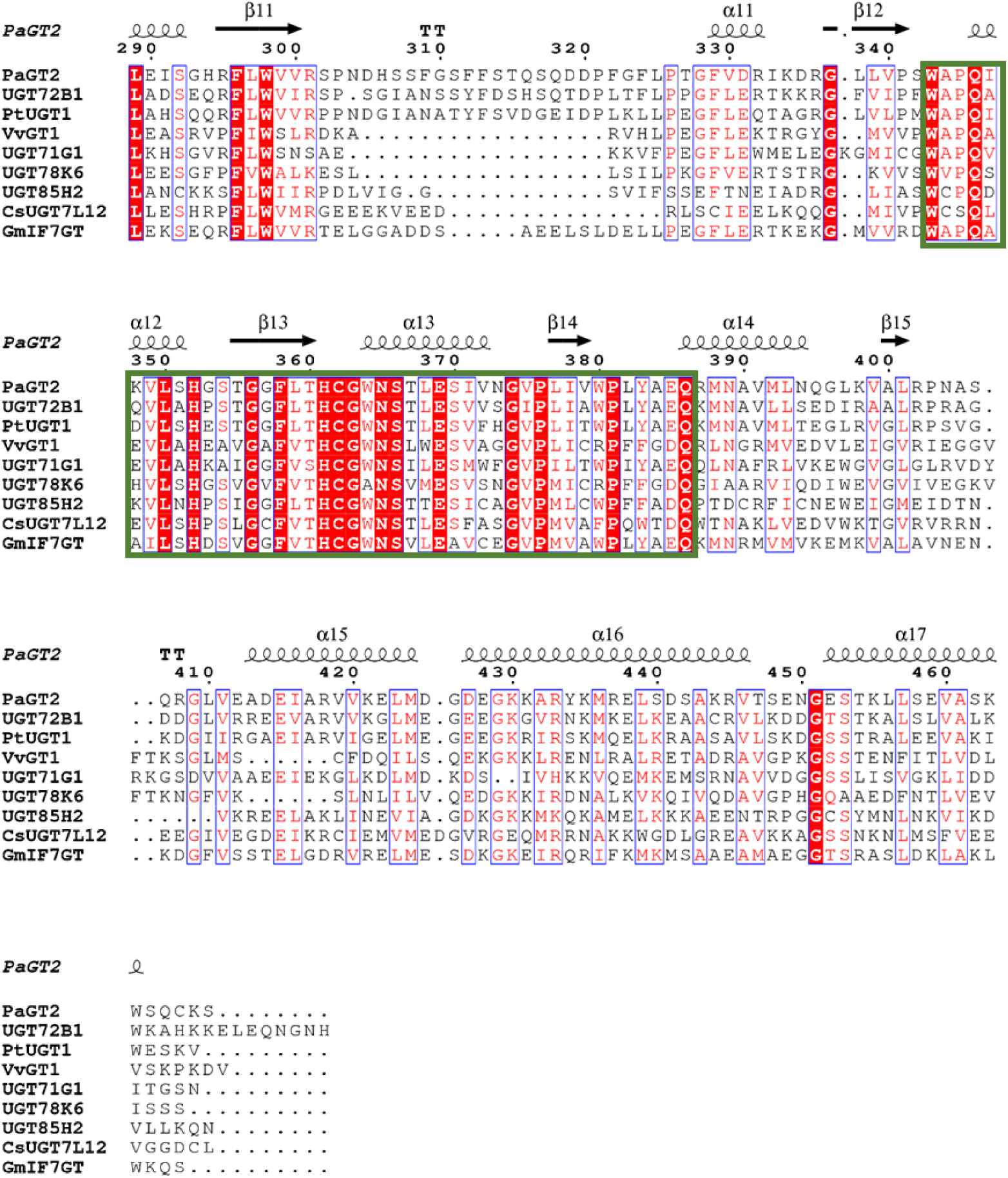
Sequence alignment of *Pa*GT2 with other UGTs. Multiple sequence alignment of *Pa*GT2 amino acid sequence with other plant glycosyltransferases UGT72B1 (*Aradopsis thaliana*), *Pt*UGT1 (*P. tinctorium*), *Vv*GT1 (*Vitis vinifera*), UGT71G1 (*Medicago truncatula*), UGT78K6 (*C. ternatea*), UGT85H2 (*Medicago truncatula*), *Cs*UGT75L12 (*Camellia sinensis*) and *Gm*IF7GT (G. max). PSPG motif is indicated by green lined box. Highly conserved residues are highlighted in red, well conserved residues are red colored. The conserved catalytic histidine (His18) is indicated with red *, the alternative catalytic histidine (His81) is indicated with green * and Cys142 is indicated with black *. This figure is drawn by using ESPript 3.0 (3)

**S11.**
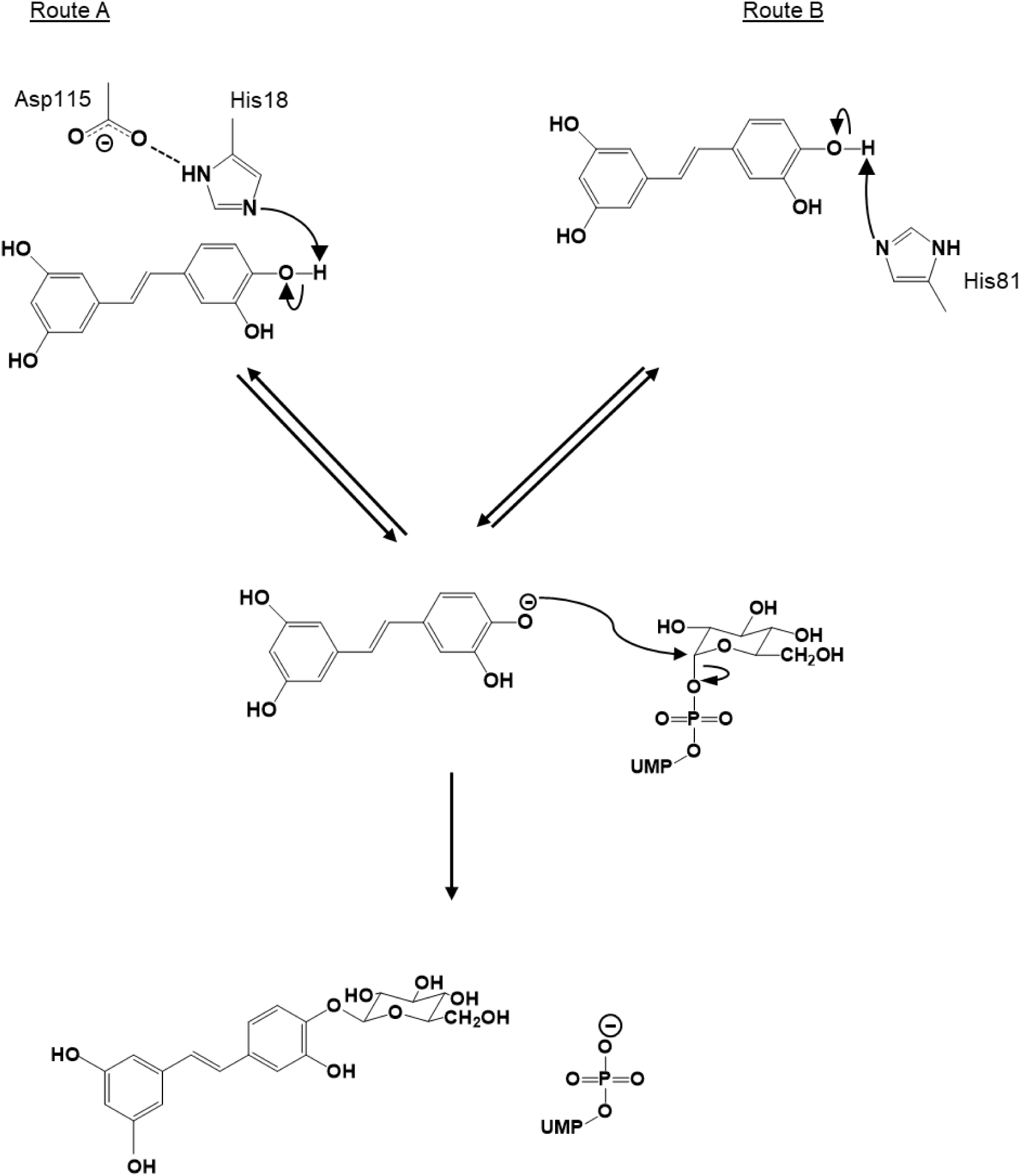
The proposed glucosylation mechanisms in *Pa*GT2. The possible mechanism of piceatannol glucosylation by *Pa*GT2. Piceatannol anion necessary for attack on C1 carbon of UDP-glucose can be generated in two different ways (A) by the catalytic pair His18 and Asp115 (B) by His81.

